# Natural killer cell cytotoxicity shapes the clonal evolution of B cell leukaemia

**DOI:** 10.1101/2023.11.16.567430

**Authors:** Michelle C. Buri, Mohamed R. Shoeb, Aleksandr Bykov, Peter Repiscak, Hayeon Baik, Alma Dupanovic, Faith O. David, Boris Kovacic, Faith Hall-Glenn, Sara Dopa, Jos Urbanus, Lisa Sippl, Susanne Stofner, Dominik Emminger, Jason Cosgrove, Dagmar Schinnerl, Anna R. Poetsch, Manfred Lehner, Xaver Koenig, Leïla Perié, Ton N. Schumacher, Dagmar Gotthardt, Florian Halbritter, Eva M. Putz

**Affiliations:** St. Anna Children’s Cancer Research Institute (CCRI), Vienna, Austria; Center for Physiology and Pharmacology, Institute of Pharmacology, Medical University of Vienna, Vienna, Austria; Center for Physiology and Pharmacology, Department of Neurophysiology and Pharmacology, Medical University of Vienna, Vienna, Austria; Division of Molecular Oncology & Immunology, Oncode Institute, Netherlands Cancer Institute, Amsterdam, The Netherlands; Christian Doppler Laboratory for Next Generation CAR T Cells, Vienna, Austria; Institut Curie, Université PSL, Sorbonne Université, CNRS UMR168, Laboratoire Physico Chimie Curie, Paris, France; Biomedical Genomics, Biotechnology Center, Center for Molecular and Cellular Bioengineering, Technische Universität Dresden, Germany; Department of Hematology, Leiden University Medical Center, Leiden, The Netherlands; Institute of Pharmacology, University of Veterinary Medicine Vienna, Vienna, Austria

**Keywords:** cancer immunoediting, natural killer cells, tumour evolution, DNA barcoding, cytotoxicity, effector function, interferon-γ, perforin, Ly6A, LY6E

## Abstract

The term cancer immunoediting describes the dual role by which the immune system can suppress and promote tumour growth and is divided into three phases: elimination, equilibrium and escape. The role of NK cells has mainly been attributed to the elimination phase. Here we show that NK cells play a role in all three phases of cancer immunoediting. Extended co-culturing of DNA barcoded mouse BCR/ABL^p185+^ B acute lymphoblastic leukaemia (B-ALL) cells with NK cells allowed for a quantitative measure of NK cell-mediated immunoediting. Whereas most tumour cell clones were efficiently eliminated by NK cells, a certain fraction of tumour cells harboured an intrinsic primary resistance. Furthermore, DNA barcoding revealed tumour cell clones with secondary resistance, which stochastically acquired resistance to NK cells. NK cell cytotoxicity put a selective pressure on B-ALL cells, which led to an outgrowth of primary and secondary resistant tumour cell clones, which were characterised by an IFN-γ signature. Besides well-known regulators of immune evasion, our analysis of NK cell resistant tumour cells revealed the upregulation of genes, including *Ly6a*, which we found to promote NK cell resistance in leukaemic cells. Translation of our findings to the human system showed that high expression of *LY6E* on tumour cells impaired the physical interaction with NK cells and led to worse prognosis in leukaemia patients. Our results demonstrate that tumour cells are actively edited by NK cells during the equilibrium phase and use different avenues to escape NK cell-mediated eradication.

## Introduction

Cancer immunoediting refers to the dynamic interaction between immune and cancer cells during tumour development (1). The immune system can have both tumour-promoting and -suppressive effects. This process occurs in three phases: elimination, equilibrium, and escape. In the elimination phase, adaptive and innate immune cells, particularly cytotoxic T and natural killer (NK) cells, recognise and eliminate cancer cells. The equilibrium phase involves the coexistence of tumour cells and the immune system, leading to sculpting of the tumour under constant immune pressure, ultimately resulting in immune escape (2).

NK cells are innate lymphoid cells with the ability of recognising and eliminating damaged, stressed, and infected cells including cancer cells (3). NK cell activation relies on a balance of activating and inhibitory signals from receptors on their surface during the interaction with potential target cells. On the one hand, NK cells kill target cells through exocytosis of cytotoxic granules containing perforin and granzymes (4), induction of death receptor-mediated apoptosis (5) or antibody-dependent cellular cytotoxicity (6). On the other hand, NK cells secrete cytokines such as interferon-γ (IFN-γ), which activate other immune cells in their vicinity (7). In the context of cancer immunoediting, NK cells play a crucial role in eliminating tumour cells (8) and limiting their metastatic spread (9). NK cells spare healthy tissue and represent a safe treatment option with minimal risk of causing adverse effects (10). As a result, there is growing interest in using NK cells for immunotherapies, which will benefit from a better understanding of how tumour cells evade NK cell attacks. Most studies to date have focused on the interaction between human NK cells and tumour cell lines obtained from patients (11–16). These tumour cells have already successfully evaded from the patient’s immune system and may thus not represent suitable models to study immunoediting.

Here, we used an *in vitro* co-culture model based on non-edited NK cell-naïve mouse BCR/ABL^p185+^ B cell acute lymphoblastic leukaemia (B-ALL) cells to investigate the primary interaction of tumour and NK cells. NK cell resistant tumour cells that emerged during the co-culture showed an upregulation of many IFN-γ-dependent genes, such as lymphocyte antigen 6 complex, locus A (*Ly6a*). This study identified Ly6A as novel driver of NK cell-mediated immune evasion of mouse leukaemia and extended our knowledge about the role of LY6E in limiting NK cell anti-tumour activity (17). Moreover, we established a model to quantify NK cell effector functions and their ability to edit tumour cells by cellular DNA barcoding (18–20). In summary, this study shows that NK cells play a role in all three phases of cancer immunoediting and contribute to the active sculpting of tumour cells, ultimately driving tumour evasion.

## Material and methods

### Mice

C57BL/6JRj wild-type mice (RRID:MGI:2670020) were obtained from the Janvier labs and C57BL/6-Prf1^tm1Sdz^/J (*Prf1^−/−^*) (21) (Strain #002407, RRID:IMSR_JAX:002407) and B6.129S7-Ifng^tm1Ts^/J (*Ifng^−/−^*) (22) mice (Strain #002287, RRID:IMSR_JAX:002287) were obtained from the Jackson Laboratory. B6.129-B2m^tm1Jae^ (*B2m^−/−^*, RRID:MGI:2175714) (23) mice were kindly provided by Thomas Kolbe (VetMedUni Vienna). The mice were between 8 and 12 weeks of age and maintained under specific pathogen-free conditions according to FELASA guidelines.

### Cell culture

Phoenix-ECO (ATCC, RRID:CVCL_H717) and Lenti-X packaging cells (Takara, RRID:CVCL_1926) were cultured in DMEM medium (Thermo Fisher Scientific) with 10% fetal bovine serum (FBS, Capricorn Scientific). The K562 cell line (ATCC, RRID:CVCL_0004) was cultured in RPMI medium (Gibco) supplemented with 100 U/ml penicillin and 100 µg/ml streptomycin (Gibco), 10% FBS and 50 µM β-mercaptoethanol (Sigma-Aldrich) (cRPMI).

To generate BCR/ABL^p185+^ (B-ALL) cell lines, BM was isolated from 8-10-week-old mice and single cell suspensions were prepared through repeated homogenisation using 18G and 24G needles. 48 hours before the infection of BM cells, Phoenix-ECO cells were transiently transfected with a retroviral MSCV-BCR/ABL^p185^ vector (kindly provided by Veronika Sexl) mixed with PureFection transfection reagent (System Biosciences). The BM cells were transduced for 1 hour with the fresh viral supernatant in the presence of 10 µg/ml polybrene (Sigma-Aldrich) and 10 ng/ml recombinant murine IL-7 (Peprotech). Transformed B-ALL cells were maintained in cRPMI medium until they were stable cell lines characterised by continuous growth and the homogenous expression of CD19, B220 and CD43 (approx. 2 months).

All cell lines were regularly tested for their mycoplasma negativity using the PCR Mycoplasma detection kit (ABM). Cell growth and survival was measured using a CASY counter (OLS OMNI Life Science) or a CellDrop BF (DeNovix).

Primary mouse NK (mNK) cells were isolated from spleens using the negative selection EasySep mouse NK cell isolation kit (STEMCELL Technologies) or positive selection CD49b (DX5) MicroBeads kit (Miltenyi Biotec) and cultured in cRPMI supplemented with 5,000 U/ml recombinant human IL-2 (rhIL-2, Miltenyi Biotec). mNK cells were identified as single live mCD45.2^+^mCD3^-^NK1.1^+^mCD335^+^ cells. Human NK (hNK) cells were isolated from heathy donors with the negative selection kit, RosetteSep Human NK Cell Enrichment Cocktail (Stemcell Technologies) and cultured in NK MACS medium (Miltenyi Biotec) supplemented with 5% human serum (Capricorn Scientific) and 500 U/ml rhIL-2. hNK cells were determined as single live hCD45^+^hCD3^-^CD56^bright^ ^or^ ^dim^ cells.

### DNA barcoding

#### Generation of the lentiviral barcode library LG2.1

The BC1DS_lib oligo containing a 21nt random barcode sequence (**Table S1**) was PCR amplified (14 cycles: 10 sec 98°C, 30 sec 57°C, 20 sec 72°C) with Phusion polymerase (New England Biolabs (NEB)). The PCR amplified product was column purified (MinElute PCR cleanup kit, Qiagen) and subsequently digested with XhoI and AscI, followed by ligation into the 3’ UTR of the GFP cDNA sequence within the pRRL lentiviral vector (24), using the Electroligase kit (NEB). Electrocompetent ElectroMax Stbl4 bacteria (Invitrogen) were electroporated with 6 ng ligation product and a small fraction of the transformed bacteria was plated on Luria-Bertani (LB) agar plates to determine transformation efficiency, while the remaining bacteria were grown overnight in 400 ml LB medium (VWR Life Science) supplemented with 100 µg/ml ampicillin (Sigma-Aldrich). DNA was extracted from the bacterial culture using the Maxiprep kit (Invitrogen). To determine the library’s barcode diversity and distribution, barcodes were amplified with a three PCR reaction protocol (see below) in biological and technical duplicates and sequenced on a HiSeq 3000/4000 device (Illumina) with 64 bp single-end reads at the Biomedical Sequencing Facility (BSF) Vienna.

#### Barcoding of tumour cells

Lenti-X cells were transfected with the LG2.1 barcode library and the two packaging plasmids p8.9QV and pVSVG (25) (provided by Ton Schumacher) using the PureFection transfection reagent. Virus supernatant was harvested after 48 and 72 hours and concentrated with AMICON columns (Sigma-Aldrich). 2×10^5^ B-ALL cells were spinfected with serially diluted viral supernatant for 90 min at 900xg. The transduction rate (%GFP^+^ cells) was measured by flow cytometry 3 days after transduction. Tumour cells with the desired transduction rate (<5% (18)) were expanded and FACS sorted to enrich GFP^+^ cells. Barcoded tumour cells were frozen for further experiments.

### Flow cytometry

Cells were analysed on a FACSymphony A3 Cell Analyzer device (Becton Dickinson, BD) and sorted on a FACSAria Fusion device (BD). The antibodies and cell stains used in this study are listed in **Table S2**. Single living cells were gated according to size and granularity in FSC-A, SSC-A and FSC-H plots and cell surface marker expression was quantified by median fluorescence intensity (MFI).

#### *In vitro* co-culture system and tumour cell growth analysis

9×10^5^ barcoded B-ALL cells were co-cultured with mNK cells (4 days after isolation) in an effector-to-target (E:T) ratio of 1:1 in three biological replicates per condition. Depending on NK cell cytotoxicity and tumour cell growth, cells were harvested for FACS sorting and further analyses (described below) on day 4-6 and 14-17. On the day of harvesting, 3×10^5^ tumour cells were co-cultured for a second or third time with 4 day old mNK cells (E:T = 1:1). All conditions were cultured in the presence of 2,500 U/ml rhIL-2. In the experiment shown in **Figures 5A&S7A**, tumour cells were treated with 2 ng/ml mouse IFN-γ (Abcam) every 2-3 days. During the co-culture, the absolute number of B-ALL tumour cells was determined by flow cytometry using AccuCheck Counting Beads (Invitrogen). Tumour cells were identified by gating on single live mCD45.2^+^mCD19^+^GFP^+^ cells.

#### Cytotoxicity assay

5×10^4^ CellTrace Violet (CTV, Invitrogen) stained B-ALL or K562 cells were seeded in 96-well round bottom plates and IL-2-activated mNK or hNK cells added in the indicated E:T ratios in technical duplicates or triplicates. After 1 or 4 hours of incubation at 37°C, cells were stained with fixable viability dye eFluor 780 (Invitrogen) and fixed with fixation buffer (Biolegend) before flow cytometric analysis. The percentage of specific NK cell lysis was calculated as follows: (% dead tumour cells after co-incubation with NK cells - % spontaneous lysis) / (100 % - % spontaneous lysis).

#### Effector function assay

5×10^4^ CTV^+^ B-ALL or K562 cells were cultured with 5×10^4^ IL-2 activated mNK or hNK cells, respectively. As positive control served mNK or hNK cells stimulated with cell stimulation cocktail (Invitrogen) or 10 ng/ml IL-12 (Peprotech) + 100 ng/ml IL-15 (Peprotech) + 15 ng/ml IL-18 (R&D System), respectively. 1 hour after the start of co-culture, Brefeldin A (Biolegend) was added, and 3 hours later cells were fixed with fixation buffer and permeabilized with intracellular staining permeabilization wash buffer (Biolegend) before the intracellular staining of the cells for IFN-γ, granzyme B and TNF-α.

#### Conjugation assay

Tumour cells were stained with CTV and NK cells were either stained with anti-mNK1.1 or anti-hCD56. An equal amount of tumour and NK cells was added into a FACS tube and cells were centrifuged for 3 min at 100xg. After no (human) or 10 min (mouse) incubation at 37°C, cells were vortexed for 3 sec and fixed with ice cold 0.5% paraformaldehyde (Electron Microscopy Sciences) in PBS. Cells were kept on ice and immediately measured by flow cytometry. Living cells were gated according to size and granularity in FSC-A and SSC-A plots, doublets were included in the analysis, and CTV and CD56 or NK1.1 double positive tumour-NK conjugates were quantified.

### Multi-omics approaches

10^6^ GFP^+^ barcoded tumour cells were FACS sorted, washed twice with PBS and snap frozen. DNA and RNA were extracted using the AllPrep DNA/RNA Mini Kit (Qiagen).

#### DNA barcode sequencing

Library preparation for barcode sequencing was performed in technical duplicates with three nested PCR steps. PCR#1 amplified the barcodes, PCR#2 added the required sequences for the universal Illumina adapters and four random base pairs (bps) for a more diverse sequencing start. PCR#3 attached the P5 and P7 universal Illumina adapters and with the P7 a sample-specific index sequence (primer sequences are in **Table S1**). The samples were cleaned up with Agencourt AMPure XP beads (Beckman Coulter). Up to 96 equimolar pooled samples spiked with 10-15% PhiX (Illumina) were sequenced with the MiSeq Reagent Kit v3 (150 cycles) (Illumina) 64 bp single-end reads with a coverage of 50-100 reads/barcode.

#### Barcode/Cell clone evolution analysis

The R statistics software v4.0.3 (RRID:SCR_001905) was used to analyse barcode abundance and clonal diversity. Unmapped reads were loaded from FASTQ format (function *readDNAStringSet* from package Biostrings v2.58.0, RRID:SCR_016949) into the R environment. Barcode sequences were located by pattern matching (function *vmatchPattern* from package Biostrings v2.58.0) to identify the fixed head (“GAACACTCGAGATCAG”) and non-variable (“TGTGGTATGATGT”) portions of the reads, which enclose the barcodes from left and right, respectively. Sequences that did not meet the expected length (21 bases) or included uncalled bases (“N”) were discarded (mean discarded sequences = 15%). To define the barcodes of the reference library (**Figure S1C-E**), we selected sequences with a sum of at least 100 reads across samples (n = 4) of the viral library for further analysis. With these we constructed a count matrix indicating the number of reads matching each barcode. Technical replicates were merged by adding up the read counts. For each cell line, only barcodes detected across three B-ALL alone samples at the corresponding reference timepoint with a total sum of at least 9 reads were considered for downstream analysis. Barcode diversity for each sample was quantified using Shannon diversity as follows *sum*(*p* ∗ *log*(*p*), *na*. *rm* = *TRUE*) ∗ −1, where *p* is the ratio of the counts of each barcode relative to the total barcode counts. We used DESeq2 (RRID:SCR_015687) v1.30.0 to find the subset of differentially abundant barcodes (absolute log_2_(fold change_cutoff) ≥ 1 & padj < 0.05) between the reference time point and conditions of interest. Shifted log_2_ transformation was used for normalisation (function *normTransform* from package DESeq2 v1.30.). Based on the statistics of differential abundance and variability across samples (here, the variability was calculated as following: *v* = *log*2(*max*(*x*+ 1)/*min*(*x* + 1)), where *x* is a vector of normalised counts across samples) at every time point, barcodes were assigned to one of five categories: i) primary resistant: differentially abundant (padj < 0.05) and increasing (log_2_(fold change) ≥ 1; comparing B-ALL + NK vs B-ALL alone); ii) eliminated: differentially abundant (padj < 0.05) and decreasing (log_2_(fold change) ≤ −1; comparing B-ALL + NK vs B-ALL alone); iii) static: variability v ≤ 0.5 in B-ALL alone, and v < 1 at B-ALL + NK; iv) secondary resistant: low variability v ≤ 0.5 in B-ALL alone, high variability in B-ALL + NK v ≥ 1 and barcode abundance high in only one of the three replicates; v) others: none of the above. Barcodes were further ordered using hierarchical clustering (function *hclust* from package *stats* v4.0.3, method = “ward.d2”, RRID:SCR_014673). Heatmaps were generated using ComplexHeatmap (RRID:SCR_017270, v2.7.5.1) and bubble plots were generated using Matlab (RRID:SCR_001622).

#### RNA sequencing

After RNA isolation (see above), samples were treated with DNase I (Thermo Fisher Scientific). Library preparation and sequencing was performed at the Biomedical Sequencing Facility (BSF) of the Research Center for Molecular Medicine (CeMM) of the Austrian Academy of Sciences. RNA-seq libraries were prepared with the QuantSeq 3‘ mRNA-Seq Library Prep Kit (FWD) for Illumina (Lexogen). For sequencing, samples were pooled into NGS libraries in equimolar amounts. Expression profiling libraries were sequenced on HiSeq 3000/4000 or Novaseq 6000 instruments (Illumina) following a 50-, 100 and 120-base-pair single-end setup. Raw data acquisition and base calling was performed on-instrument, while the subsequent raw data processing off the instruments involved two custom programs based on Picard tools (RRID:SCR_006525, v2.19.2). In a first step, base calls were converted into lane-specific, multiplexed, unaligned BAM files suitable for long-term archival (IlluminaBasecallsToMultiplexSam, 2.19.2-CeMM). In a second step, archive BAM files were demultiplexed into sample-specific, unaligned BAM files (IlluminaSamDemux, 2.19.2-CeMM).

#### RNA-seq analysis

Mapping of the reads was performed using STAR (26) (RRID:SCR_004463, v2.7.10a) to the *Mus musculus* GRCm38, or *Homo sapiens* GRCh38 assemblies and duplicate reads were identified with Picard (v2.27.4). Quantification of the mapped reads was carried out using Salmon (27) (RRID:SCR_017036, v1.8.0). The following data processing and visualisation was performed in R (v4.2.0). To ensure robustness of the analysis, the data set was filtered to include only transcripts detected in at least three samples. The analyses of mouse cell lines A and B and cell lines C and D were performed separately. Finally, the results were compared.

For count normalisation, variance stabilisation, and differential expression analysis, the Bioconductor DESeq2 (28) (v1.38.3) package was used. DESeq2 employs a negative binomial distribution-based model, with additional shrinkage using apeglm (29) or ashr algorithm (v1.20.0). Transcripts were considered significantly differentially expressed between groups of comparison if they exhibited a log_2_(fold change) ≥ 0.58 and a padj ≤ 0.05.

The principal component analysis (PCA), of expression heatmaps and volcano plots were generated using ggplot2 (v3.4.2) and heatmap (v1.0.12). To correct for batch effects between different experiments in PCA plots (**Figures S3A&B, S4A**) the limma R (30) package (RRID:SCR_010943, v3.54.2) was utilised. The resulting gene lists were annotated and subjected to gene set enrichment analysis (GSEA, RRID:SCR_003199). Visualisation of the GSEA results was achieved using the hypeR (31) package (v2.0.1) and represented as dot plot. Gene Ontology Biological Process (GOBP) terms were obtained from the Molecular Signatures Database (MSigDB) using the msigdbr (32) package (RRID:SCR_016863, v7.5.1).

#### ATAC sequencing

5×10^4^ GFP^+^ tumour cells were FACS sorted into 1.5 ml micro centrifuge tubes containing 400 µl 0.5% Bovine Serum Albumin (BSA, GE Healthcare) in PBS and pelleted. Cell lysis and DNA tagmentation was performed in 25 µl tagment DNA enzyme in tagment DNA buffer (Illumina), 1% digitonin (Promega) and EDTA-free protease inhibitor cocktail (Roche) for 30 min at 37°C. Samples were purified with the Qiagen MinElute kit (Qiagen). Libraries were generated as described by Buenrostro et al (33) by using the Ad1_noMx and Ad2.1-Ad2.48 indexed primers (**Table S1**). PCR products were purified with Agencourt AMPure XP beads and the sample profile was analysed with the High Sensitivity D5000 kit (Agilent Technologies) on a TapeStation (Agilent Technologies). 45-48 equimolar pooled samples were sequenced on a NovaSeq 6000 instrument with 50 bp paired-end reads at the BSF (CeMM).

#### ATAC-seq analysis

The obtained raw reads were processed using the ATAC-seq Next Flow (34) pipeline (v2.0). This included mapping the reads with STAR to the GRCm38 genome, identification of duplicates with Picard (v2.27.4), calling broad peaks with MACS2 (35) (v2.2.7.1). Peaks overlapping with blacklisted (GRCh38) regions were filtered out and the remaining peaks were annotated with HOMER (36) (RRID:SCR_010881, v4.11). Consensus peaks across all samples were identified with BEDTools (37) (RRID:SCR_006646, v2.30.0) and counting reads in these peaks was done with featureCounts (38) (RRID:SCR_012919, Rsubread v2.0.1). Visualisation of the ATAC-seq peaks was performed by creation of BigWig files with bedGraphToBigWig (v377) and exporting to the IGV (39) browser (RRID:SCR_011793, v2.16.1). Quality of the obtained raw data was assessed with MultiQC (40) (RRID:SCR_014982, v1.13). Data processing and visualisation was performed in R (v4.2.0). To ensure robustness of the analysis, the data set was filtered to include only consensus peaks detected in all three replicates of each condition and cell line and only the peaks that were located in the window of ±3kb around the transcription start site (TSS). The analysis of cell lines A and B was performed separately from the analysis of cell line C and D.

Similar to the RNA-seq analysis, the Bioconductor DESeq2 (1.38.3) package was used for count normalisation, variance stabilisation, and analysis of differential accessibility. Peaks were considered significantly accessible between groups of comparison if they exhibited a log_2_(fold change) ≥ 0.58 and a padj ≤ 0.05. Exploratory analysis included PCA, production of heatmaps, volcano plots and GSEA.

#### Integration of RNA-seq and ATAC-seq data

To integrate two datasets and perform genome-wide quantification of differential transcription factor activity in cell lines at different time points we used diffTF (41) (v1.8). The processing was done only for A and B data sets, due to very high computational demand of the algorithm. The input consisted of raw ATAC-seq reads in BAM files, filtered consensus peaks from the previous step, raw RNA-seq counts, and PWM data for all TFs – HOCOMOCO (42) (RRID:SCR_005409, v10).

#### 2D-RP/RP Liquid Chromatography – Tandem Mass Spectrometry analysis

Cells were lysed in 200 μL lysis buffer (50 mM HEPES pH 8.0, 2% SDS, 1 mM PMSF, cOmplete Protease Inhibitor Cocktail, Roche), heated to 99 °C for 5 min, and DNA was sheared using a Covaris S2 ultrasonicator. Lysates were cleared at 20,000 × g for 15 min at 20 °C and tryptic digest was performed using FASP according to Wisniewski et al (43). Peptides were desalted (C18 SPE; Pierce peptide desalting columns, Thermo Fisher Scientific), labelled with TMTpro™ 16plex reagents (Thermo Fisher Scientific), pooled, dried in a vacuum concentrator, and cleaned up via C18 SPE. After re-buffering in 10 mM ammonium formiate buffer pH 10, peptides were fractionated via a C18 column (150 × 2.0 mm Gemini-NX, 3 µm, C18 110Å, Phenomenex, Torrance, CA, USA) on a Dionex UltiMate 3000 nanoLC (Thermo Fisher Scientific). Fractions were dried in a vacuum concentrator and taken up in 0.1% TFA before analysis.

Mass spectrometry analysis was performed on an Orbitrap Fusion Lumos Tribrid mass spectrometer coupled to a Dionex UltiMate 3000 nanoLC system via a Nanospray Flex Ion Source (all from Thermo Fisher Scientific) interface using the same procedure as Block et al (44). Data analysis was performed using the Proteome Discoverer platform (RRID:SCR_014477, v.2.4.1.15).

### Genome editing

#### Ribonucleoprotein (RNP)-mediated CRISPR genome editing of B-ALL cells

Cells were transfected with crRNAs, Alt-R CRISPR Cas9 tracrRNA ATTO 550 and Alt-R S.p. Cas9-GFP V3 (IDT) with the Gene Pulser Xcell electroporation system (BioRad) using the following settings: 350 V, 5 ms and 1 pulse. crRNA sequences: *Ly6a*: TCACGTTGACCTTAGTACCC; TATTGAAAGTATGGAGATCC, *Plaat3*: CGTCATGTTTGTTATTGACC; ATGGCCCAGTGTCTGTACAT. The next day, GFP^+^ATTO 550^+^ tumour cells were FACS sorted. Cells were single cell FACS sorted into 96-well flat bottom plates into 50% tumour cell conditioned medium and 50% fresh cRPMI. After the outgrowth of single clones, the KO was validated by TIDE sequencing. Up to 2×10^5^ cells were lysed in 50 µl Pawe’s buffer at 56°C for 15 min with subsequent heat inactivation at 95°C for 5 min. After centrifugation at 2000xg for 5 min, supernatant was used for the TIDE PCR to amplify the breakpoints of the guide RNAs (primer sequences in **Table S1**). PCR products were Sanger sequenced (Microsynth) and analysed for their genotype with the <u>TIDE (nki.nl)</u> web tool (RRID:SCR_023704). Gene knockouts were validated using flow cytometry or quantitative PCR.

#### CRISPR genome editing in Cas9 expressing K562 cell line

K562 were transduced to express Cas9 with lentiviral supernatant from Lenti-X cells 48 hours after transfection with the following constructs: lentiCRISPRv2 (RRID:Addgene_52961), p8.9QV and VSV-G (both plasmids provided by T. Schumacher). K562 cells were transduced by spinfection for 90 min at 900xg. Transduced cells were selected with 1 µg/ml puromycin (Sigma-Aldrich) for up to 7 days. crRNA specific for LY6E (GTGACTGTGTCTGCTAGTGC) was transfected into K562-Cas9 cells with the Gene Pulser Xcell electroporation system using following settings: 400 V, 3 ms and 1 pulse. The cells were single cell FACS sorted into 96-well flat bottom plates into 50% 48 hours old medium of tumour cells and 50% fresh cRPMI with 100 Gray irradiated K562-mbIl15-41-BBL (kindly provided by Manfred Lehner). After the outgrowth of single clones, the KO was validated by TIDE sequencing (primer sequences in **Table S1**) and immunoblotting.

#### Quantitative PCR

To validate the *Plaat3*-deficiency in B-ALL cells, RNA was extracted from 2×10^6^ cells with the Monarch Total RNA Miniprep Kit with “on column” DNase I treatment (NEB). cDNA was generated by using LunaScript RT SuperMix Kit (NEB) and qPCR was performed with the Luna universal qPCR Master Mix (NEB) (primer sequences in **Table S1**). The qPCR was conducted on a 7500 Real Time PCR System device (Applied Biosciences) and the data was analysed with the 7500 Software v2.0.6.

#### Immunoblotting

To analyse LY6E protein expression, *LY6E* WT and KO cells were cultured with or without 10,000 U/ml IFN-β (Miltenyi Biotec) or 10 ng/ml IFN-γ (Peprotech) overnight. Cells were lysed in RIPA buffer (Sigma-Aldrich) with protease inhibitor cocktail (Thermo Scientific) and phosphatase inhibitor cocktail II (Abcam). Equal amounts of protein were separated on a SDS polyacrylamide gel. Proteins were transferred to a nitrocellulose membrane by using the Trans-Blot turbo transfer kit and system (Bio-Rad). After blocking in 5% BSA-TBS-T, membranes were probed with primary antibodies against LY6E (Invitrogen, RRID:AB_3075213) and β-actin (Cell Signaling Technologies, RRID:AB_2223172) overnight, before the staining with a fluorescent secondary antibody (Thermo Scientific, RRID:AB_614946) and visualisation by the ChemiDoc MP imaging system (Bio-Rad).

### Seahorse – ATP rate assay

3.5 ×10^4^ *Ly6a* WT or KO cells were seeded in triplicates into the Seahorse XF HS PDL Miniplates (Agilent) and centrifuged at 300xg for 2 min. The assay was conducted according to manufacturer’s protocol, measured on the Agilent Seahorse XF HS Mini Analyzer (Agilent) and analysed with the online Agilent Seahorse Analytics software.

### IFN-γ ELISA

Tumour cells were split to 3×10^5^ cells/ml on day 0 and cultivated for 48 hours in cRPMI. On day 2, the supernatant was harvested, filtered through a 45 µm filter, and transferred onto IL-2 expanded mNK cells in the presence of 2,500 U/ml rhIL-2. After 48 hours, mNK cells were harvested and the supernatant was frozen at −80°C. As positive control served the stimulation of mNK cells with cell stimulation cocktail 4 hours before the supernatant harvesting. The ELISA was performed according to manufactures protocol (Abcam) and analysed using the analysis tool: “https://www.arigobio.com/elisa-analysis”.

### Statistical analysis

Statistical analysis was performed using GraphPad Prism (RRID:SCR_002798, v8.4.3 for Windows) using two-tailed paired and unpaired t-tests, one-way, two-way ANOVA or Kruskal-Wallis tests as indicated. Where ANOVA showed a statistical difference, Tukey’s multiple comparison testing or Dunnett’s post-hoc testing was applied. The α level for all tests was set to 0.05, and p values were two tailed. The significance level is indicated as following: *p < 0.05; **p < 0.01; ***p < 0.001; ****p < 0.0001.

To generate the heatmaps in Figure 1B&S1A, the data was pooled from 2-4 independent experiments by calculating the mean surface marker expression or mean % specific lysis at E:T=10:1. The values were standardised for each variable (row) using the following calculation: Z-score = (mean of observed values – mean of the samples) / (standard deviation of the samples).

**Figure 1:**
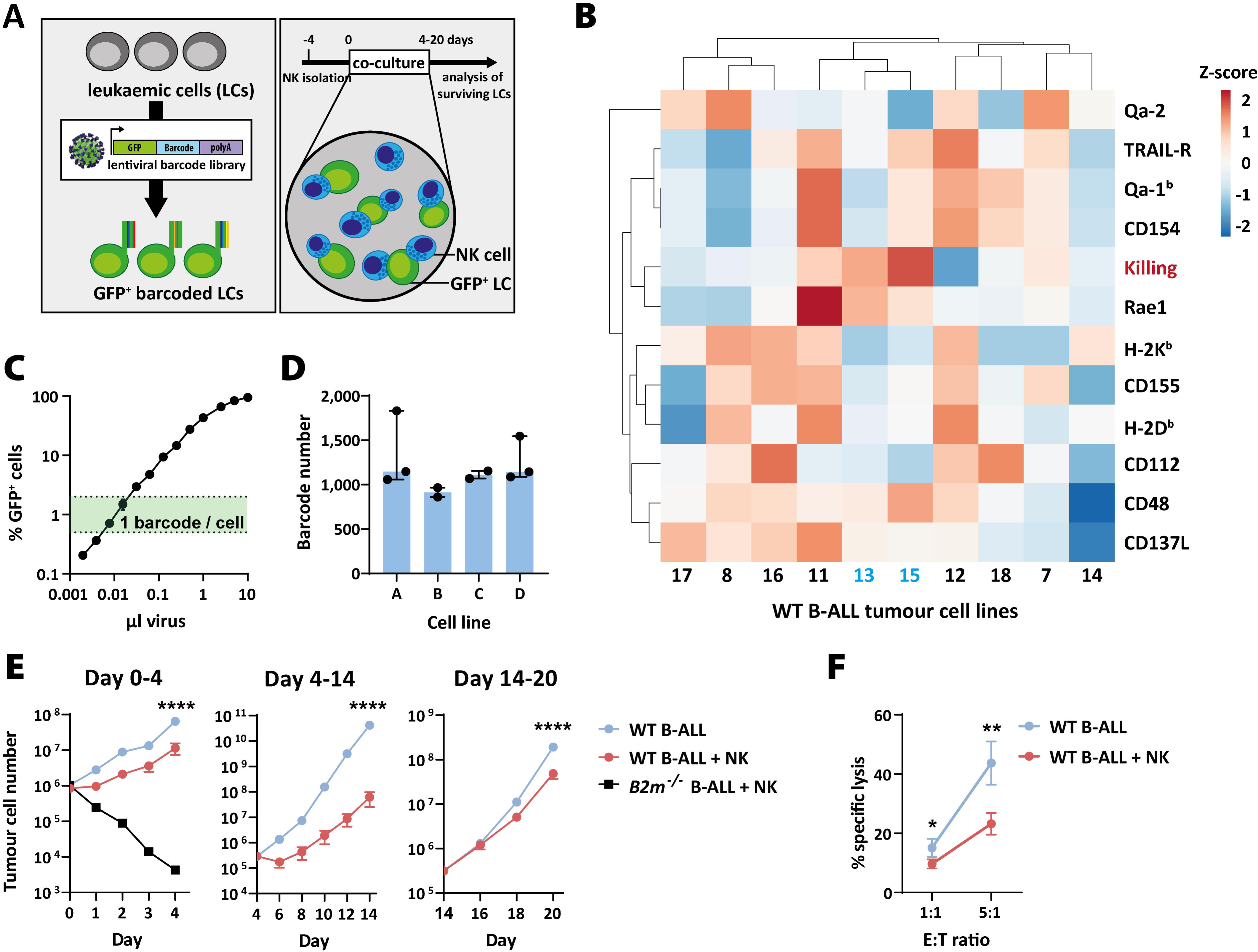
Co-culture system to study cancer immunoediting *in vitro*. **(A)** Scheme illustrates the experimental setup: BCR/ABL^p185+^ B-ALL leukaemic cells (LCs) are transduced with a DNA barcode library. Barcoded LCs are co-cultured with NK cells for 4-20 days *in vitro*. **(B)** Ten B-ALL cell lines were generated from C57BL/6 WT mice and characterised regarding the surface expression of NK cell receptor ligands (black) and their susceptibility towards NK cell killing (red, depicted here is the cytotoxicity at E:T=10:1) measured by flow cytometry. Cell lines #13 and #15, in blue, showed the highest NK cell susceptibility and were used for the following experiments. Rows were centred; unit variance scaling was applied to rows. Both rows and columns were clustered using correlation distance and average linkage. The plot was generated using the Clustvis web tool (80). **(C)** B-ALL cell lines were DNA barcoded by lentiviral transduction. Using different amounts of viral supernatant, the optimum transduction efficiency of <5% GFP^+^ cells (18) was chosen to allow a single barcode integration per cell. Shown are means±SD (n=2) of one representative cell line. **(D)** Two B-ALL cell lines (#13 and #15) were barcoded and the barcode diversity of the FACS-purified GFP^+^ cell lines was determined by targeted DNA sequencing. Cell lines A and B were derived from #13 and cell lines C and D from #15. Bars show barcode diversity of barcoded tumour cell lines (A&D n=3 and B&C n=2 independent experiments, each determined by sequencing of 3 biological and 2 technical replicates). Shown is median with interquartile range. **(E)** Barcoded WT B-ALL cell lines A-D were cultivated in three individual wells either in the absence or presence of purified and IL-2-activated NK cells. One *B2m^−/−^*B-ALL cell line was included as control. Absolute numbers of B-ALL cells were determined by flow cytometry. On day 0, 4 and 14, B-ALL cells were FACS sorted, re-plated and fresh NK cells were replenished. Shown are means±SD of n=4 B-ALL cell lines of one representative experiment (n=3 independent experiments); the significance was calculated by an unpaired t-test on day 4, 14 and 20. **(F)** On day 29, a 4 hour NK cytotoxicity assay was performed using B-ALL cells that had been cultured for 20 days in the presence or absence of NK cells. Shown are means±SD of n=4 cell lines (in technical triplicates) of one representative experiment (n=3 experiments); the significance was calculated by a paired t-test.

Heatmaps in Figures 5I&S7F were generated by using the R package ComplexHeatmap (v2.10.0). The raw median fluorescence intensity (MFI) values minus the MFI of their unstained control served as input. Rows (surface markers) were scaled and clustered (unsupervised). Columns (cell lines) were ordered by genotype and cell line.

### Data Availability

The data generated in this study are publicly available in Gene Expression Omnibus (GEO) at <u>GSE278011</u> (DNA barcode sequencing), <u>GSE278010</u> (ATAC sequencing) and <u>GSE278006</u> (RNA sequencing). The computer codes for the data analysis are available on Github (https://github.com/TumorImmunoEditingLab).

## Results

### Quantitative measure of NK cell effector functions

As a model to study NK cell evasion of leukaemic tumour cells we established a long-term *in vitro* co-culture system using IL-2-activated primary mouse NK cells co-incubated with NK cell-naïve DNA barcoded B-ALL cell lines for up to 20 days (**Figure 1A**). We generated several independent B-ALL cell lines from the bone marrow (BM) of wild-type (WT, n=10) and major histocompatibility complex 1 (MHC-I)-deficient mice (*B2m^−/−^*, n=10) and characterised their NK cell receptor ligand surface expression and susceptibility towards NK cells. The two WT cell lines, which showed the highest susceptibility towards NK cells (#13 and #15), expressed enhanced levels of the activating NKG2D-ligand Rae1 and lower levels of inhibitory ligands (e.g. classical and non-classical MHC-I proteins H2-K^b^, H2-D^b^, Qa-1^b^ and Qa-2) (**Figure 1B**). *B2m*^−/−^ B-ALL cell lines showed a higher susceptibility towards NK cells according to the recognition of missing self (45) (**Figure S1A**). To allow for a systematic quantification of NK cell effector functions, a DNA barcode library was introduced into the WT B-ALL cell lines #13 and #15 (n=2 each) to track individual tumour cell clones over time. For this purpose, a new lentiviral barcode library (LG2.1, containing 52,645 barcodes) was generated consisting of 21 random nucleotides positioned downstream of a green fluorescent protein (*GFP*) gene (**Figures S1B-E**). The lentiviral library was introduced into the B-ALL cell lines at a low multiplicity of infection to ensure the integration of a single barcode per cell (25,46) (**Figure 1C**). To enrich for barcoded cells, GFP^+^ cells were FACS sorted shortly after transduction and before the long-term co-culture with NK cells. The barcode diversity of the resulting four cell lines called A and B (derived from #13), C and D (derived from #15) was determined by site-specific PCR amplification and next generation sequencing (**Figure 1D**). The barcoded cells were co-incubated with NK cells at an effector to target (E:T) ratio of 1:1. After 4 and 14 days, GFP^+^ tumour cells were FACS purified and fresh NK cells were replenished to re-challenge the tumour cells. Throughout the experiment, the absolute number of GFP^+^ tumour cells was determined by flow cytometric analysis (**Figure 1E**). While the tumour cells initially showed significantly slower proliferation in the presence of NK cells during the first two rounds of co-culture, this difference became marginal in the third round. Twenty-nine days after the start of co-culture, we performed a cytotoxicity assay. The extended co-culture of leukaemic cells with NK cells induced resistance towards NK cell-mediated killing and led to tumour evasion *in vitro* (**Figure 1F**).

### NK cell-mediated editing of leukaemic cells

There are four conceivable fates of B-ALL tumour cell clones upon long-term co-culture with NK cells: The abundance of one clone could be lower (“eliminated”), higher (“primary resistant”), unchanged (“static”) or highly variable (“secondary resistant”) (**Figure 2A**). After bulk DNA sequencing of the barcodes, we designed a decision tree to discriminate these four categories algorithmically (**Figure 2B**). Primary resistance, also known as intrinsic resistance (47), is defined as *a priori* resistance towards therapy. In our study, primary resistance was characterised by a reproducible and significant accumulation of a clone despite the presence of NK cells in co-culture. In comparison, secondary resistance describes the phenomenon of acquired resistance, represented by a situation in which a tumour can initially respond effectively to therapy but relapse or progress after treatment (48). We hypothesised that secondary resistance was not a general tumour-intrinsic property but acquired only by a subfraction of sibling cells and would therefore be observed only in isolated wells (**Figure 2A**). Indeed, clustering of the barcoded leukaemic cells according to the pre-defined categories (**Figures 2C&D**), showed that the majority of clones was efficiently eliminated (58.5±4.8%, n=2-3 independent experiments, n=2-4 B-ALL cell lines, mean±SD). A smaller number of cell clones showed an NK cell resistant phenotype (14.3±4%) or remained static (23.9±4.4%) despite the presence of NK cells. As hypothesised, we also observed secondary resistance, characterised by the presence of a cell clone in only one of the three replicate wells (3.3±2.5%, **Figure 2E**). In line with the paradigm that the acquisition of a resistant phenotype in tumour cells takes time, we observed the emergence of secondary resistance primarily at the later of the two sampling timepoints (**Figure S2A**). To be able to draw conclusions about the pre-existing fate of the tumour cell clones, we compared two independent experiments with the same barcoded cell lines. As shown in **Figures 2F and S2B**, the cell clones attributed to the clusters primary resistant and eliminated had the same fate in both experiments. In contrast, the pool of secondary resistant tumour cell clones was distinctive for independent experiments. In summary, whereas primary resistant and eliminated cells harboured tumour-intrinsic properties *a priori* that determined their destiny, the fate of secondary resistant tumour cell clones was stochastic. This underscored our hypothesis that secondary resistant tumour cells acquired their resistant phenotype during NK cell co-culture. Our data clearly showed that apart from direct killing, NK cells participated in immunoediting by actively shaping the tumourigenicity of individual tumour cells.

**Figure 2:**
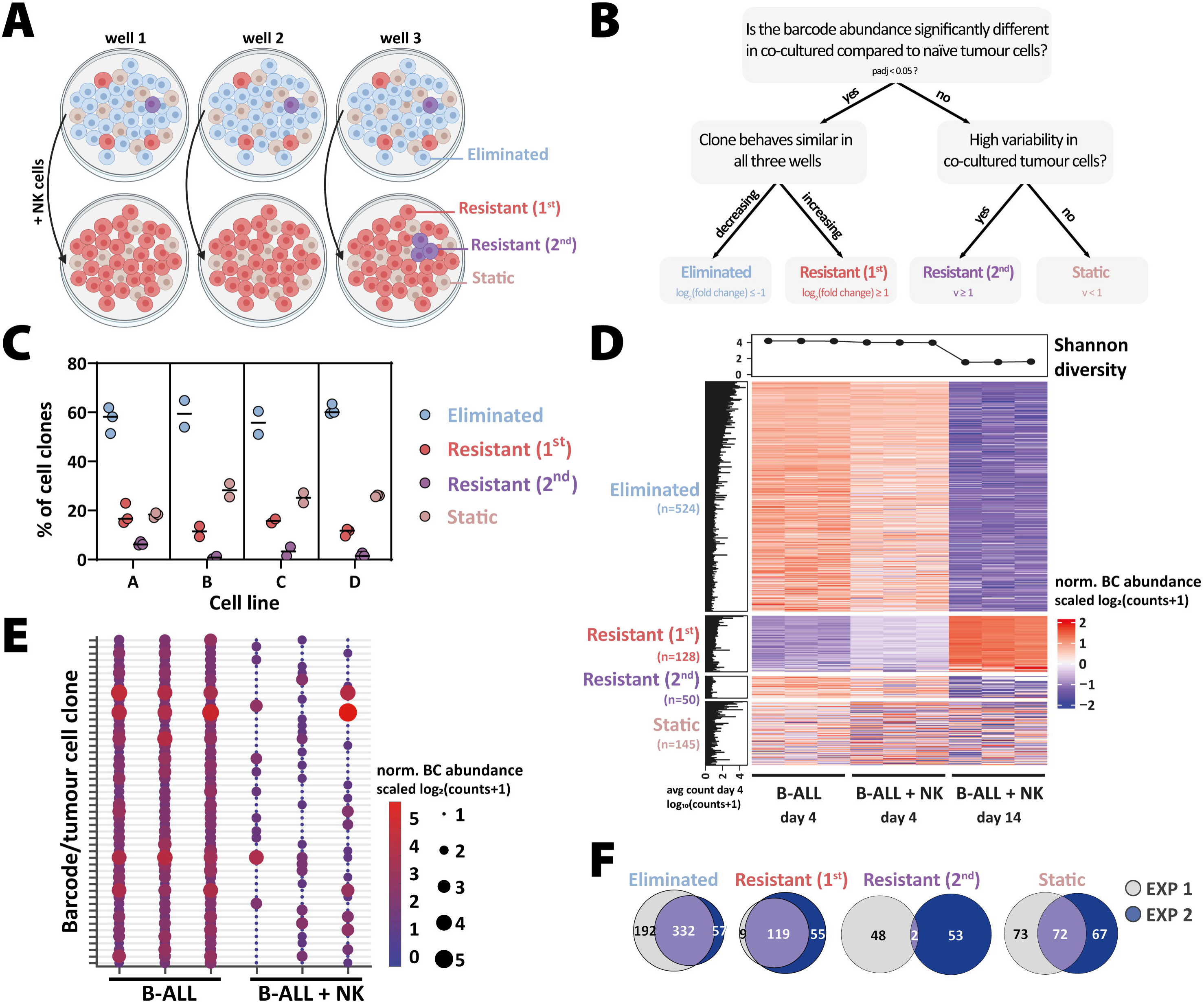
Quantification of NK cell-mediated cancer immunoediting *in vitro*. **(A)** Schematic representation of the proposed model. Scheme was created in BioRender. Kovar, H. (2023) BioRender.com/j11m245. **(B)** Upon long-term co-culture of B-ALL and NK cells, we hypothesised that each tumour cell clone would fall into one of the following categories: The abundance of a clone can be significantly higher (primary resistant) or lower (eliminated), unchanged (static) or show a high variability (secondary resistant) upon NK cell co-culture. Variability is defined as *v* = *log*2(*max*(*x*+ 1)/*min*(*x*+ 1)), where *x* is a vector of normalised counts across samples. A rule-based decision tree was designed to discriminate the four categories depicted in (A). **(C)** The percentage of the cell clones in each group on day 14 is depicted for cell lines A, B, C and D. Each data point represents the mean value (n=3 wells, sequenced in duplicates) of n=2-3 independent experiments. **(D)** The heatmap shows the normalised abundance of barcodes (norm. BC abundance) of cell line A from one representative experiment. Each subcolumn (n=3) in the figure represents a sample (well), and each row represents a barcode. The barcodes were divided into the four pre-defined groups according to the criteria defined in (B). The Shannon diversity index (shown above the heatmap) serves as measure of barcode diversity and drops significantly after 14 days of NK co-culture. The histogram on the left shows the average abundance of the barcoded cell clone in the B-ALL only samples on day 4. **(E)** The bubble plot depicts the normalised abundance of secondary resistant clones in B-ALL samples (day 4) compared to B-ALL + NK samples (day 14). The x-axis shows the three individual wells of each condition, while each row on the y-axis shows an individual tumour cell clone. The size and colour of the bubbles indicate the normalised barcode abundance. Shown is the same cell line and experiment as in (D). **(F)** Comparing two independent experiments in cell line A as shown in (D&E), the Euler diagrams highlight a high overlap of eliminated, primary resistant, and static cell clones. In contrast, only two secondary resistant clones were shared between both independent experiments.

### Molecular changes observed in NK cell resistant B-ALL cells

To identify novel drivers of NK cell resistance we investigated changes in the transcriptomic and chromatin accessibility profiles of NK cell resistant B-ALL cells generated during the long-term *in vitro* co-culture. To this end, we performed RNA (**Figures 3A-D and S3**) and ATAC sequencing (**Figures 3E&F and S4**) and compared NK cell co-cultured to NK cell naïve tumour cells. The transcriptomic analysis showed a significant upregulation of genes with a well-known link to NK cell immune evasion such as components of the MHC-I machinery (e.g., *Stat1*, *B2m*, *Tap1* and *H2-K1* (49)) (**Figures 3A and S3C**). Additionally, this analysis revealed a novel set of genes, which may include potential drivers of NK cell resistance, such as *Ly6a* and phospholipase A and acyltransferase 3 (*Plaat3*). The functional enrichment analysis showed that the main biological process associated with the differentially expressed genes (DEGs) was the response to IFN-γ (e.g., *Stat1*, *Irf1*, *Ccl5* and *Cd74*) (**Figure 3B**), which is produced by activated NK cells in co-culture. After integrating the results from the four barcoded cell lines A-D, we observed that many DEGs upon NK cell co-culture were cell line specific (up: A/B=193 and C/D=99, down: A/B: 211 and C/D=63). However, a total of 129 genes were up- and 27 were downregulated in all four cell lines and may be universally associated with or driving NK cell escape (**Figure 3C**). The top 30 DEGs are depicted in the heat map in **Figure 3D**. In line, the ATAC sequencing results showed the opening of chromatin at the promoters of classical IFN-inducible genes (e.g., *Iigp1*, *Ifi27*, *Mx1*), as well as of *Ly6a* and *Plaat3* (**Figures 3E and S4C**). We considered overlapping differentially accessible regions (DARs) in cell lines A/B and C/D as the most robust ones (**Figure 3F**). Integration of RNA and ATAC sequencing data by comparing the DEGs and DARs in cell lines A/B showed a significant intersection (37 genes up/open, 16 genes down/closed). The two highly upregulated genes *Ly6a* and *Plaat3* in the transcriptomic analysis were also present in this strictly filtered data set (**Figure 3G**). The quantification of differential transcription factor (TF) activity by integrating the RNA and ATAC sequencing data sets revealed TFs involved in NK cell resistance. We classified TFs as activators (NF2L1, SMRC1, JUN, IRF1 and IRF9) or repressors (RXRB, JUND), based on their expression and accessibility of predicted binding sites (50). STAT1 expression was clearly upregulated but showed activating and repressive functions and was therefore classified as undetermined (**Figure 3H**). A closer look at the top hits *Ly6a* and *Plaat3* showed that both have multiple predicted binding sites for STAT1, IRF1 and IRF9 based on the occurrence of DNA sequence motifs in the vicinity of accessible chromatin (**Figure 3I**). In summary, multiomics analyses revealed that the B-ALL cells, which were not *a priori* sensitive and eliminated by NK cells, showed a highly NK cell resistant phenotype characterised by a strong IFN-γ signature and the upregulation of *Ly6a* and *Plaat3*.

**Figure 3:**
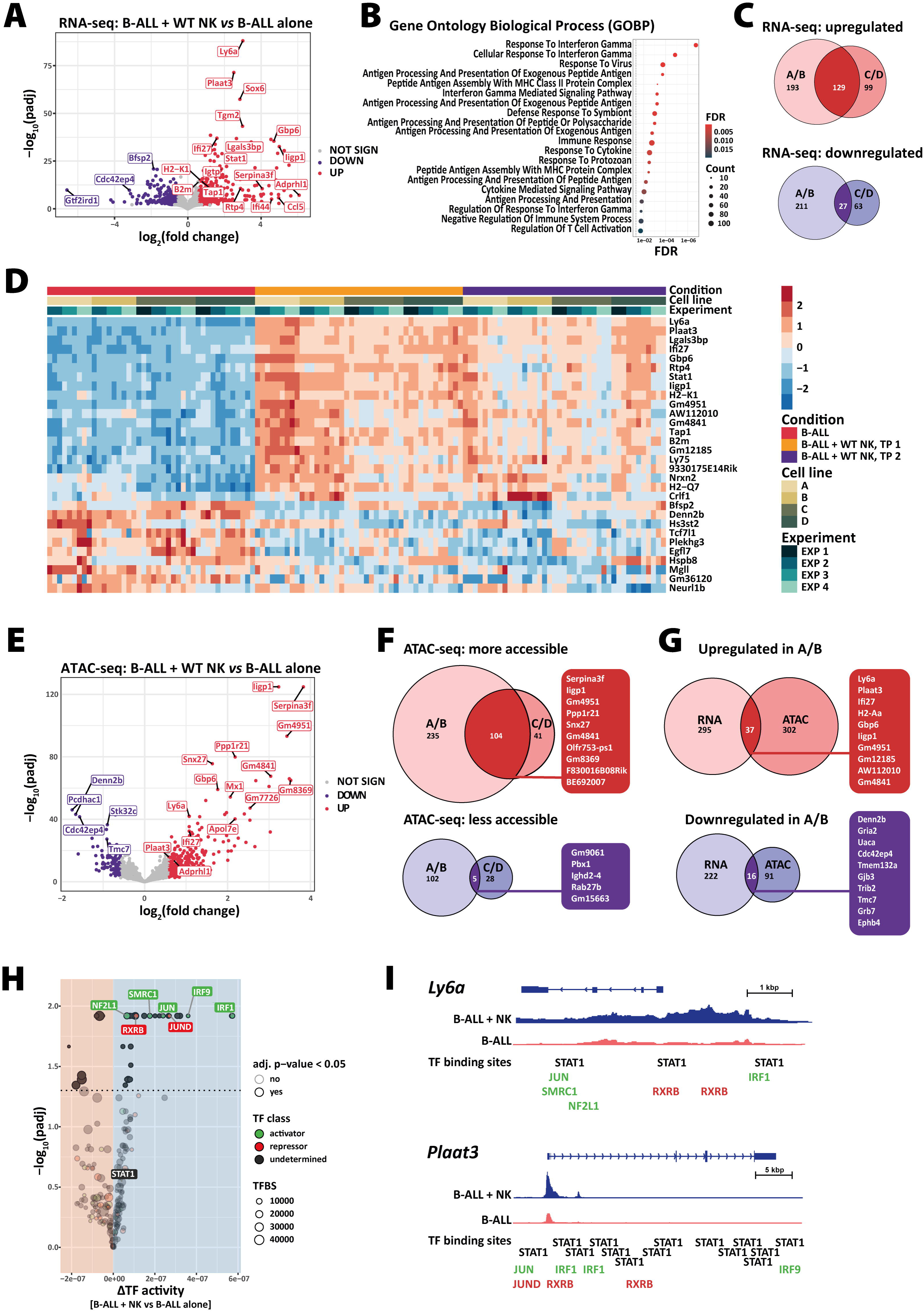
Integrative analysis of RNA and ATAC sequencing shows well-known and novel genes deregulated in NK cell resistant B-ALL cells. The **(A-D and G)** transcriptome and **(E-I)** chromatin accessibility analysis of B-ALL cells co-cultured with NK cells for 4 (time point 1 (TP1)) or 14 (TP2) days were analysed and compared to the B-ALL alone cells on TP1 (n=3-4 independent experiments). **(A)** The volcano plot shows the significantly up- or downregulated genes in B-ALL cell lines A and B after co-culture with NK cells for 14 days (n=3 experiments). Statistics were calculated with the Wald test. **(B)** The dot plot illustrates the enrichment analysis using the hyperR tool (31) for the Gene Ontology Biological Process (GOBP) terms associated with NK cell co-cultured B-ALL cells shown in panel (A). X-axis and dot colour depicts the false discovery rate (FDR) and y-axis the corresponding biological processes. Dot size illustrates the number of genes associated with the biological processes. **(C)** The Euler diagrams display up- or downregulated genes in cell lines A/B and/or C/D after 14 days of NK cell co-culture. **(D)** The heatmap shows the expression of selected overlapping genes (top-20 upregulated, top-10 downregulated; ranked by p-value) identified in (C) (n=3-4 experiments, 4 B-ALL cell lines). The row-scaled normalised counts represent the log_2_(fold change). **(E)** The volcano plot shows DARs in B-ALL cells A/B after co-culture with NK cells for 14 days (n=3 experiments). Statistics were calculated with the Wald test. **(F)** The Euler diagrams highlight genes whose promoters (TSS ± 3kb) had higher or lower chromatin accessibility in cell lines A/B and/or C/D after 14 days of NK cell co-culture in n=3-4 experiments. Gene promoters with the highest log_2_(fold change) are highlighted. **(G)** Integration of RNA and ATAC sequencing data shows that 37 genes are up- and 16 downregulated in cell lines A/B; the top 10 up- or downregulated genes are highlighted on the right. **(H)** Analysis of differential transcription factors activity using (diffTF) (41) highlighted activator (green) and repressor (red) TFs that were differentially expressed in B-ALL cell lines A/B after NK cell co-culture. **(I)** Integrative Genomics Viewer (IGV) genome browser tracks display representative ATAC sequencing signal densities at the *Ly6a* and *Plaat3* loci in B-ALL cells with or without NK cell co-culture. Shown are the gene bodies including a 3 kb upstream promoter region, as well as binding sites of TFs identified in (H). Database HOCOMOCO (v10) (42) was used and depicted are counts per million (CPM) with a range of [0-1.37].

### Relative contribution of NK cell cytotoxicity and IFN-γ production in cancer immunoediting

The two main functions of NK cells are the direct lysis of tumour cells and the production of pro-apoptotic and -inflammatory cytokines, most importantly IFN-γ. To discriminate the relative contributions of killing and cytokine production to cancer immunoediting, we performed two independent long-term co-culture experiments with NK cells deficient in perforin (*Prf1^−/−^*), which renders them unable to kill their target cell (21), or deficient in IFN-γ production (*Ifng^−/−^*) (**Figures 4, S5 and S6**). As described above, the growth of GFP^+^ tumour cells was measured by flow cytometry during the experiment. On day 4 and 14 (for *Prf1^−/−^)* or on day 4 and 17 (for *Ifng^−/−^*), GFP^+^ tumour cells were sorted and co-cultured with freshly prepared NK cells. As previously observed, B-ALL cells co-cultured with WT NK cells showed an initial susceptibility during the first two weeks and developed an NK resistance thereafter (**Figures 4A&C**). Co-culture with *Prf1^−/−^* NK cells only led to a minor growth impairment of the tumour cells (**Figure 4A**). The presence of *Ifng^−/−^*NK cells diminished the number of B-ALL cells in the beginning of the co-culture, but the tumour cells became resistant sooner than B-ALL cells co-cultured with WT NK cells (**Figure 4C**). Five days after B-ALL cells outcompeted NK cells, we measured their NK cell resistance. A cytotoxicity assay using WT NK cells was performed and showed that in both experiments, B-ALL cells that had been co-cultured with WT NK cells showed the highest NK cell resistance. *Prf1^−/−^* as well as *Ifng^−/−^* NK cell co-cultured B-ALL cells were more resistant compared to B-ALL cells cultured alone, however the resistance was lower than in WT NK co-cultured B-ALL cells (**Figures 4B&D**).

**Figure 4:**
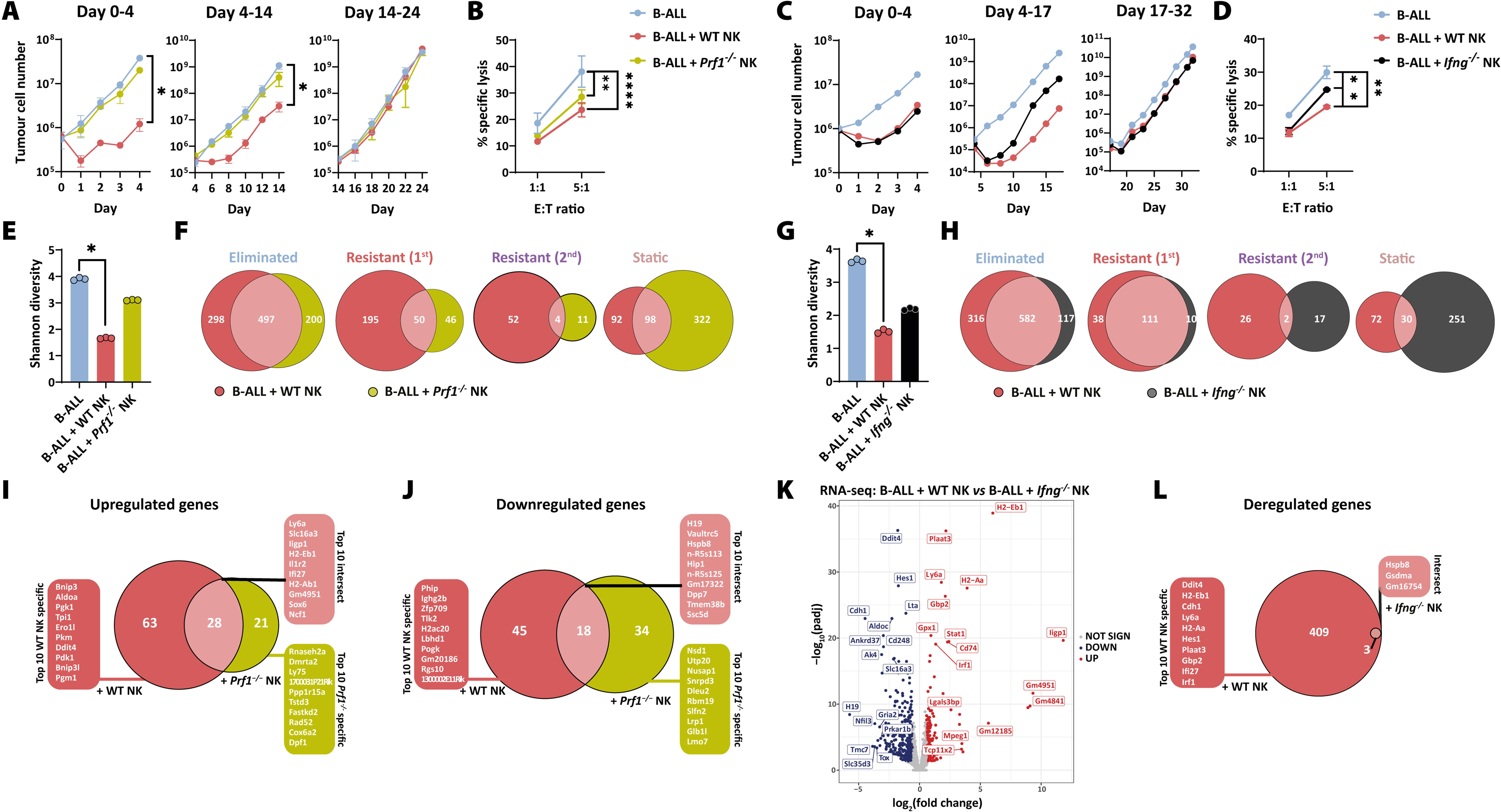
The combination of NK cell cytotoxicity and IFN-γ production drives tumour immunoediting. **(A&C)** Barcoded B-ALL cell lines (n=2 cell lines A/D) were cultivated in three individual wells either in the absence or presence of purified and IL-2-activated WT, **(A)** *Prf1^−/−^* or **(C)** *Ifng^−/−^* NK cells. Absolute numbers of B-ALL cells were determined by flow cytometry. On day 0, 4 and 14/17 B-ALL cells were FACS sorted, re-plated and fresh NK cells were replenished. **(A)** Shown are means±SD of n=2 cell lines. The significance was calculated with the Kruskal-Wallis test and Dunńs multiple comparisons testing. **(B&D)** On the last day of the NK cell co-culture experiment, a standard 4 hour NK cytotoxicity assay with WT NK cells was performed using B-ALL cells that had been cultured for 4 weeks in the presence or absence of NK cells. Shown are means±SD of technical triplicates. Statistics were calculated for the highest E:T ratio with one-way ANOVA with Tukeýs multiple comparisons test. **(E&G)** The Shannon diversity of leukaemic clones dropped significantly (*p=0.0219*) upon co-culture with WT NK cells, and to a lesser degree with **(E)** *Prf1^−/−^*or **(G)** *Ifng^−/−^* NK cells. Shown is the mean and dots depict each replicate. Statistics were calculated with the Kruskal-Wallis and Dunńs multiple comparisons testing. **(F&H)** The barcodes within each sample were classified according to the decision tree in Fig. 2B. The barcodes that were attributed to the four pre-defined groups were compared between the treatment groups with Euler diagrams for each group [eliminated, resistant (1^st^), resistant (2^nd^) and static]. **(I&J)** The Euler diagrams display the number of up- and downregulated genes in cell lines A/D after 14 days of co-culture with WT or *Prf1^−/−^* NK cells. Top 10 up- or downregulated genes in the intersection or specific for WT or *Prf1^−/−^* NK cell co-culture are depicted in the corresponding lists. **(K)** The volcano plot depicts DEGs in B-ALL + WT NK *versus* B-ALL + *Ifng^−/−^* NK conditions on day 17 of cell line A. **(L)** The Euler diagram depicts DEGs when comparing B-ALL + WT or B-ALL + *Ifng^−/−^* NK conditions with B-ALL alone. Top 10 WT NK cell specific genes and the overlapping genes are depicted in the corresponding lists. Shown is **(A, B, I, J)** one experiment including n=2 cell lines A/D or **(C-H, K, L)** one experiment from cell line A. Data is representative for two independent experiments and n=2 cell lines.

To study the clonal behaviour of the tumour cells, the B-ALL cells were analysed for their barcode diversity on day 4 and 14/17. In line with our previous experiments, the barcode diversity of the B-ALL + WT NK cell condition dropped significantly in both experiments by approx. 58%. The Shannon diversity of B-ALL cells was decreased by 20% and 40% after co-culture with *Prf1^−/−^*or *Ifng^−/−^* NK cells, respectively (**Figures 4E&G**). We observed a strong overlap of eliminated tumour cell clones in all treatment groups, suggesting that these cell clones were *a priori* susceptible to NK cells. In line with fewer eliminated cell clones, we detected more static B-ALL clones upon co-culture with *Prf1^−/−^* or *Ifng^-/-^* NK cells. The number of primary and secondary resistant tumour cell clones was similar after co-culture with WT and *Ifng^−/−^* NK cells (**Figures 4H and S5A**). In contrast, the presence of *Prf1^−/−^* NK cells led to a drastically lower number of primary and secondary resistant tumour cell clones (**Figures 4F and S5B**). This suggested that the strong immunological pressure of NK cells leading to the emergence of primary or secondary resistance was driven by cytotoxicity rather than IFN-γ production.

To define the molecular changes driving NK cell resistance, we compared the transcriptomic profiles of B-ALL cells after co-culture with WT and *Prf1^−/−^* (day 14, **Figures 4I&J and S6A&B**) or *Ifng^−/−^* NK cells (day 17, **Figures 4K&L and S6C&D** and day 32, **Figures S6E-H**). B-ALL cells after culture with WT or *Prf1^−/−^* NK cells showed an overlap of 28 up- and 18 downregulated DEGs, including *Ly6a* and *Plaat3* (**Figures 4I&J and S6A&B**). Intriguingly, B-ALL cells only showed marginal changes in their transcriptome when cultured with *Ifng^−/−^* NK cells (**Figures 4K&L and S6C-H**). Accordingly, only B-ALL cells cultured in the presence of IFN-γ-producing WT NK cells induced the expression of NK cell receptor ligands, such as components of the classical and non-classical MHC-I machinery (**Figures S6I&J**).

In conclusion, while NK cell cytotoxicity was pivotal for the induction of primary and secondary resistance in tumour cells, IFN-γ production induced an NK cell resistant phenotype on the transcriptomic level. The combination of NK cell cytotoxicity and IFN-γ production was needed for selecting and sculpting resistant tumour cells and ultimately for tumour immunoediting.

### Role of PLAAT3 and Ly6A/LY6E in tumour evasion

NK cell resistance in B-ALL cells was associated with altered expression of well-known and novel genes involved in cancer immunoediting. We demonstrated in multiple experiments that *Plaat3* and *Ly6a* belonged to the most significantly upregulated genes in NK cell resistant B-ALL cell lines. Therefore, we followed up on these genes and studied their role in immune evasion. We performed a co-culture experiment as described in Figure 4A and analysed the B-ALL cells on day 14 by mass spectrometry to confirm the upregulation of PLAAT3 and Ly6A on the protein level. Indeed, PLAAT3 and Ly6A were increased after WT or *Prf1^−/−^* NK cell co-culture and even more upon recombinant IFN-γ treatment (**Figures 5A and S7A**). We confirmed that the IFN-γ-dependent upregulation of Ly6A in B-ALL cells in NK cell co-culture was stable, as it persisted several days after the NK cells were gone from the co-culture (**Figure 5B**).

**Figure 5:**
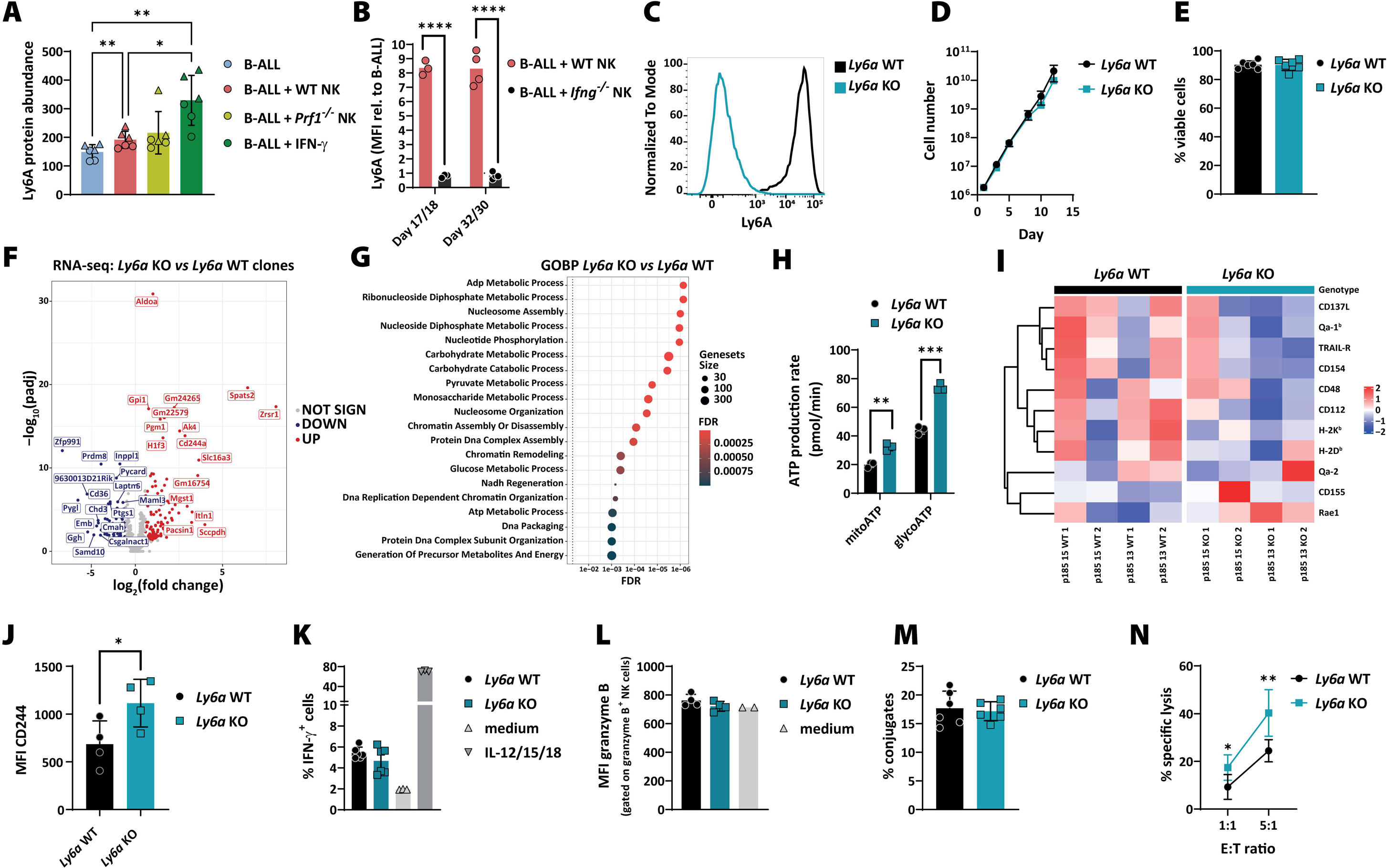
The role of mouse Ly6A in the evasion of leukaemic cells from NK cell-mediated surveillance. **(A)** The bar graph shows the protein abundance of Ly6A on day 14 of the same co-culture experiments as described in Figure 4A. Two B-ALL cell lines (A in circles/D in triangles) were analysed by mass spectrometry in triplicates. Bars and error bars represent means of arbitrary units (AU) relative to the whole protein abundance ±SD; the significance was calculated by one-way ANOVA with Tukeýs multiple comparisons test. **(B)** Ly6A surface expression was measured on B-ALL cells co-cultured with WT or *Ifng^−/−^* NK cells for 17/18 or 32/30 days. Bars represent means (n=3-4 for cell line D from two independent experiments). **(C)** *Ly6a* KO clones were generated by CRISPR/Cas9 genome editing in parental B-ALL cell lines. The gene modification was verified by measuring Ly6A surface expression by flow cytometry. Histogram shows the Ly6A expression in one representative *Ly6a* WT and KO clone. **(D)** Growth curve of *Ly6a* WT and KO B-ALL clones in absolute cell numbers over 13 days. Shown are means±SD of n=2 cell lines per genotype, measured in technical duplicates. **(E)** Cell viability of *Ly6a* WT and KO B-ALL clones. Shown are means±SD of n=2 clones per genotype of n=3 independent experiments. **(F)** The volcano plot shows DEGs (n=342) of *Ly6a* WT and KO clones (n=2 clones per genotype, sequenced in technical triplicates) under normal culturing conditions. Differentially up- or downregulated genes are depicted in red and blue, respectively. **(G)** The dot plot illustrates the GOBP terms associated with *Ly6a*-deficiency in B-ALL cells shown in panel (F). X-axis and dot colour depicts the false discovery rate (FDR) and y-axis the corresponding biological processes. Dot size illustrates the number of genes associated with the biological processes. **(H)** An ATP rate assay was performed in *Ly6a* WT and KO cells. Bar graph depicts means±SD of mitochondrial and glycolytic ATP production; the significance was calculated by an unpaired t-test. **(I)** *Ly6a* WT and KO B-ALL cell lines (n=4 per genotype) were characterised by their surface expression of NK cell receptor ligands by flow cytometry. Columns represent different tumour cell clones ordered according to genotype and cell line. Data has been scaled and clustered by row to highlight variations in marker expression across different samples. Red indicates higher and blue lower expression levels of surface ligands. **(J)** The expression of CD244 on the surface of WT and *Ly6a* KO clones was determined by flow cytometry. The bars and error bars represent the mean±SD of the MFIs of n=2 clones per genotype and two independent experiments. Statistics were calculated using an unpaired t-test. **(K)** Measurement of IFN-γ^+^ mNK cells upon co-culture with *Ly6a* WT and KO B-ALL cell lines for 4 hours (n=2 clones per genotype and technical triplicates). As positive control mNK cells were activated with IL-12/15/18. **(L)** Measurements of MFI of granzyme B in freshly isolated NK cells co-cultured with *Ly6a* WT and KO clones for 4 hours. Shown are means±SD of n=2 clones per genotype and two independent experiments. **(M)** *Ly6a* WT and KO B-ALL cell lines were co-incubated with WT NK cells for 10 min before the flow cytometric-based assessment of tumour-NK conjugate formation (n=6 clones per genotype). **(N)** A 4 hour NK cytotoxicity assay was performed using WT and *Ly6a* KO B-ALL clones. Shown are means±SD of n=2 clones per genotype and 3 independent experiments; statistics were calculated using an unpaired t-test.

To study the role of PLAAT3 in more detail, we generated *Plaat3* KO B-ALL clones and verified the lack of *Plaat3* mRNA by qPCR (**Figure S7B**). *Plaat3* KO cells did not show any difference in proliferation and viability, IFN-γ production by NK cells and NK cell receptor ligand expression (**Figures S7C-F**). Moreover, loss of *Plaat3* did not affect direct NK cell-mediated killing of mouse B-ALL cells (**Figure S7G**). To test the relevance of the human orthologue *PLAAT3* in leukaemia, we analysed the *PLAAT3* expression of ALL patients and observed an insignificant trend of worse overall survival in patients with increased *PLAAT3* expression (**Figure S7H**). These results suggested that while PLAAT3 expression was upregulated by IFN-γ and correlated with NK cell resistance, it does not constitute a driver of NK cell resistance.

Likewise, we deleted *Ly6a* in B-ALL cells and verified the lack of Ly6A surface protein expression by flow cytometry (**Figure 5C**). Ly6A deficiency had no impact on B-ALL cell proliferation or survival (**Figures 5D&E**). To investigate the impact of Ly6A on the transcriptomic level, we performed RNA sequencing of *Ly6a* WT and KO B-ALL cells and found 127 up- and 80 downregulated genes mainly related to metabolic processes (**Figures 5F&G**). In line with these results, we observed an increased mitochondrial and glycolytic ATP production in *Ly6a* KO cells (**Figure 5H**). Increased tumour metabolism may induce stress ligand expression and consequently lead to better activation of NK cells (51). While the majority of classical NK cell receptor ligands showed unaltered expression, we observed a slightly higher expression of Rae1 (*p=0.08,* unpaired *t*-test) and lower expression of CD112 (*p=0.04*) in B-ALL cells lacking Ly6A (**Figure 5I).** Further, CD244 was upregulated in *Ly6a* KO cells on RNA and protein level (**Figures 5F&J**), a marker known to be present on NK cells (52), but only basally expressed in mouse marginal-zone B cells (53) with a reported function in B cell differentiation and autoimmunity (54). We tested the functional consequences of *Ly6a* deficiency for the interaction of B-ALL and NK cells. We detected unaltered IFN-γ (**Figure 5K**) and granzyme B production (**Figure 5L**) upon co-culture with B-ALL cells and similar numbers of tumour-NK conjugates (**Figure 5M**) irrespective of the presence or absence of *Ly6a*. However, *Ly6a* KO B-ALL cells were more susceptible to NK cells compared to WT cells in cytotoxicity assays (**Figure 5N**). These results clearly show that Ly6A plays an important role in NK cell-mediated immune evasion of mouse B-ALL cells.

LY6E has been well-described as close relative of Ly6A in the human system (55). High *LY6E* expression correlated significantly with poor survival in ALL patients (**Figure 6A**). To further study the human relevance of our findings, we generated *LY6E*-deficient BCR/ABL1^+^ K562 cells and verified the lack of LY6E protein expression by immunoblotting. Notwithstanding that others have seen an IFN-β and IFN-γ-dependent induction of LY6E expression in hematopoietic cells (55), we only detected an upregulation of LY6E using IFN-β (**Figure 6B**). Transcriptomic analysis of *LY6E* WT and KO K562 cells showed a total of 829 deregulated genes (454 up and 375 down, **Figure 6C**), which were mainly associated to cell death, cell cycle and intracellular transport (**Figure 6D**). However, we did not observe any significant differences in cell proliferation nor viability between *LY6E* WT and KO clones (**Figures 6E&F**). When analysing the interaction of NK cells with *LY6E* WT and KO clones we found no difference in NK cell IFN-γ and TNF-α production (**Figure S8**), but a tendency for an increased NK cell susceptibility of the *LY6E* KO clones (**Figure 6G**). More strikingly, we detected that *LY6E* KO clones formed significantly more tumour-NK conjugates compared to *LY6E* WT cells (**Figure 6H**).

**Figure 6:**
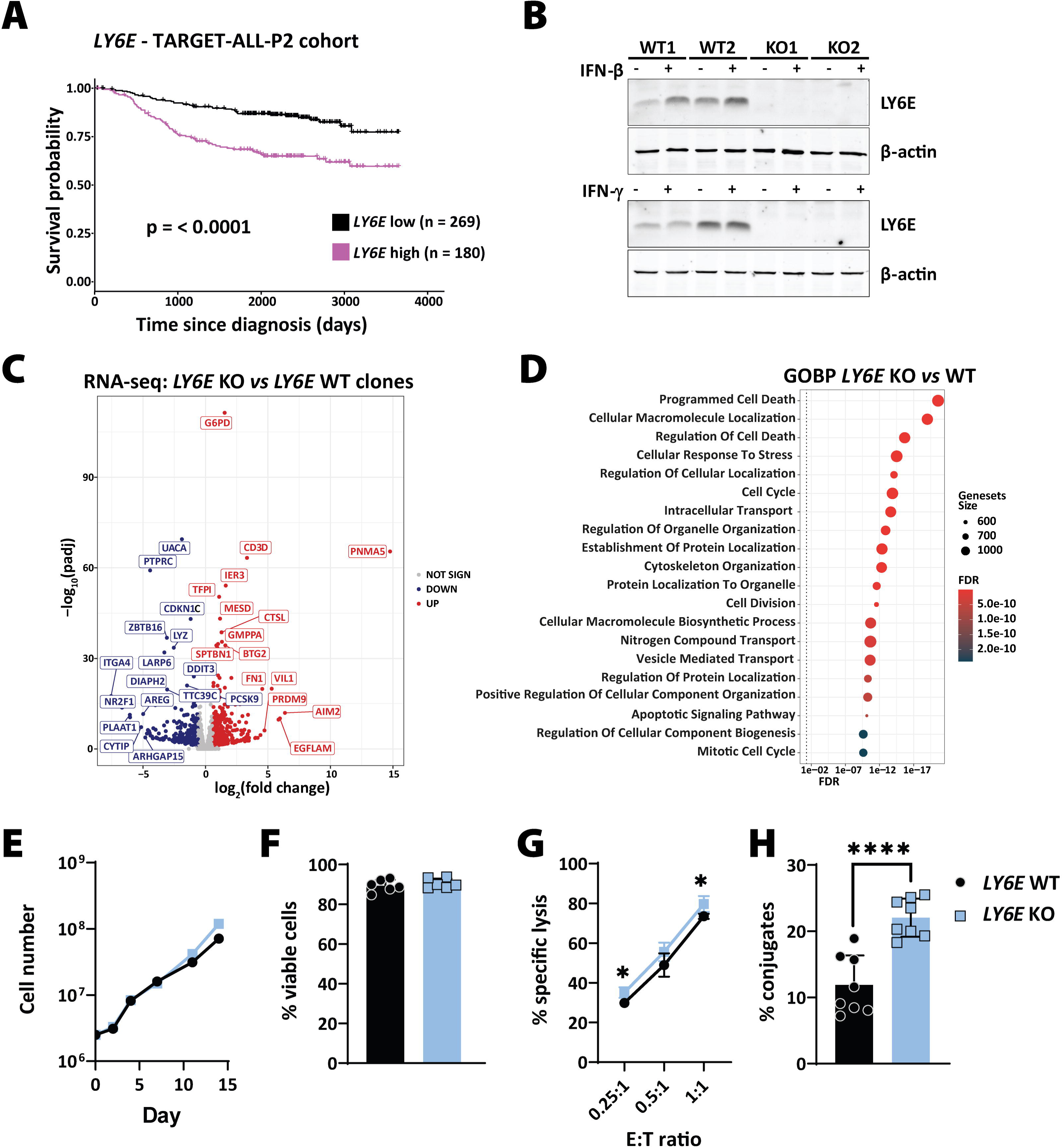
The role of human LY6E in the evasion of leukaemic cells from NK cell-mediated surveillance. **(A)** Kaplan-Meier plot depicts survival probabilities of the TARGET-ALL-P2 patient cohort divided into *LY6E* high and low expressing groups (cut-off percentile = 60%). Graph was generated by using the online web tool: <u>cSurvival (ubc.ca)</u>. **(B)** *LY6E* WT and KO K562 clones were analysed for their expression of LY6E in the absence or presence of IFN-β (upper panel) or IFN-γ (lower panel) by Western blotting. β-actin served as loading control. **(C)** The volcano plot shows DEGs (n=829) of *LY6E* WT and KO K562 clones under normal culturing conditions (n=2 clones per genotype, sequenced in technical triplicates). Differentially up- or downregulated genes are depicted in red and blue, respectively. **(D)** The dot plot illustrates the GOBP terms associated with *LY6E*-deficiency in K562 cells shown in panel (C). X-axis and dot colour depicts the false discovery rate (FDR) and y-axis the corresponding biological processes. Dot size illustrates the number of genes associated with the biological processes. **(E)** Growth curve of *LY6E* WT and KO K562 clones in absolute cell numbers over 14 days. Shown is one cell line per genotype, representative of two cell lines. **(F)** Cell viability of *LY6E* WT and KO K562 clones. Shown are means±SD of n=2 cell lines per genotype and 3 independent experiments. **(G)** A 1 hour NK cytotoxicity assay was performed using WT and *LY6E* KO K562 clones. Shown are means±SD of n=2 clones per genotype and 2 independent experiments. Statistics were calculated using an unpaired t-test. **(H)** *LY6E* WT and KO cell clones were co-incubated with NK cells and the tumour-NK cell conjugate formation was assessed by flow cytometry. Bars and error bars represent means±SD of n=2 clones per genotype and 4 independent experiments. Statistics were calculated using an unpaired t-test.

In summary, our functional analysis of PLAAT3 and Ly6A showed that their expression is upregulated in tumour cells co-cultured with NK cells in an IFN-γ-dependent manner. Although PLAAT3 was persistently upregulated in our multiomics analysis, it proved to have no effect on NK cell resistance. In contrast, Ly6A/LY6E were identified as a novel drivers of tumour evasion by impairing the interaction of NK and tumour cells.

## Discussion

NK cells represent a potent arm of the immune system, tasked with eliminating malignant cells. Nonetheless, they can fail controlling cancer cells which successfully evaded from the immune surveillance. The presented study aimed to elucidate the mechanisms employed by tumour cells to escape NK cells. Our study is based on heterogeneous, non-edited and immune-naïve mouse BCR/ABL^p185+^ B-ALL tumour cell lines, which were co-cultured for several weeks with mouse NK cells. This unique model enables to explore the primary interplay between tumour cells and the immune system. Furthermore, the B-ALL tumour cells were DNA barcoded, a technique that allows for the quantification of tumour cell clonal dynamics and a better understanding of NK cell effector functions. To the best of our knowledge, this particular aspect has not been demonstrated in previous studies. The determination of tumour cell fate during the co-culture revealed that most tumour cell clones possessed inherent sensitivity or resistance to NK cells. This technique has already been used by many others to quantify tumour cell susceptibility towards immune cells, drug- or antibody-based immunotherapies (56–59). Similar to our research, these studies observed that the majority of tumour cells is predestined for being sensitive or resistant to treatment. Our study highlighted tumour cell clones beyond those two fundamental fates of life and death, which acquired secondary resistance in the presence of NK cells. Secondary resistance, defined as the occurrence of resistance in one of three replicate wells after two weeks of co-culture, was only observed in a small subset of tumour cell clones. Similarly, using DNA barcoding to investigate the clonal evolution of drug-treated lung cancer cells, Acar *et al.* recently described the emergence of “*de novo* resistant lineages” in one replicate only (56). Interestingly, we found that secondary resistant tumour cells were highly distinctive for each independently performed experiment, suggesting that a stochastic event causes the resistance. However, due to the necessity of expanding the barcoded cell lines before the experiments (to ensure reproducible representation of all barcodes in every well), we cannot formally exclude the possibility of pre-existing heterogeneity within a cell clone. Furthermore, the barcode sequence of the LG2.1 library was found to be located approximately 900 bps upstream of the polyA tail and could thus not be detected using single-cell RNA sequencing. It would certainly be interesting to delve deeper into secondary resistance mechanisms and to compare the transcriptomic profiles of primary and secondary resistant tumour cell clones in the future (60,61).

In summary, the DNA barcode analysis demonstrated that NK cells can shape tumour cells and contribute to immunoediting of tumour cell clones. To dissect the two main functions of NK cells, namely killing tumour cells and producing IFN-γ, we analysed the clonal evolution of B-ALL cells co-cultured with *Prf1^−/−^* and *Ifng^−/−^* NK cells, respectively. It is widely acknowledged that IFN-γ is a major player in cancer immunoediting (62). In our experiments, tumour cells responded to the NK cell co-culture with an upregulation of many IFN-γ inducible genes, such as MHC-I molecules, thereby inhibiting NK cells and inducing resistance. Interestingly, and against our expectations, we observed a similar number of primary and secondary resistant B-ALL cell clones upon co-culture with WT or *Ifng^−/−^*NK cells. On the contrary, co-culture with *Prf1^−/−^* NK cells led to fewer primary and secondary resistant B-ALL cell clones. This clearly suggested that the selection of *a priori* resistant tumour cell clones and the acquisition of secondary resistance was mainly mediated by NK cell killing rather than IFN-γ production. However, the transcriptomic changes observed in long-term co-cultured B-ALL cells were only partially affected by loss of NK killing capacity, but completely absent in B-ALL cells cultured with *Ifng^−/−^* NK cells. This data argues against a major contribution of other NK cell-derived cytokines such as TNF-α on the tumour immunoediting process. We thus concluded that while IFN-γ plays only a minor role in tumour cell elimination, it enhances NK cell resistance of the remaining tumour cells by upregulating IFN-γ dependent genes, such as genes involved in MHC-I presentation and *Ly6a*.

Amongst others, previously described NK cell tumour evasion mechanisms include the upregulation of proteins involved in antigen presentation and MHC-I and -II molecules (63,64), the loss or shedding of NKG2D ligands such as MHC class I chain-related molecule A and B (MICA/B) (65) and secretion of TGF-β and IL-10 (66). Notably, we here describe *Ly6a* overexpression as emerging novel driver of tumour evasion by impairing NK cell-mediated killing. Ly6A, also called stem cell antigen-1 (Sca-1), is an 18-kDa mouse glycosylphosphatidylinositol-anchored cell surface protein. It plays an important role in haematopoietic progenitor/stem cell lineage fate, brain blood barrier transport and cancer stem cell biology (67–70). Ly6A was shown to be upregulated in cancer stem cells and many tumour entities and is induced by Wnt/β-Catenin signalling and TGF-β deregulation. This promotes cell adhesion and migration *in vitro,* increased tumourigenicity and resistance to chemotherapy *in vivo* (70–73). The human LY6 gene family comprises several members, but the direct orthologue of mouse *Ly6a* has been under debate for a long time. *LY6A/LY6S* was just recently discovered probably due to its genomic localisation on the opposite strand of and highly overlapping with *C8orf3/LINC02904* (55,74). The expression of *LY6A/LY6S* seems to be restricted to pituitary tumours (74) and a subset of non-classical lymphoid cells of the spleen (55), and it could not be detected in hematopoietic cell lines (74) or cancer patients. Besides *LY6A/LY6S*, other *LY6* family members were suggested as close homologues of the murine *Ly6a* gene, such as *LY6D*, *LY6E*, *LY6H* and *LY6K*, which were implicated as novel biomarkers for poor cancer prognosis and are frequently amplified in human cancer (75). Al Hossiny *et al.* showed that human LY6E promotes breast cancer *in vivo* and drives drug sensitivity and epithelial-to-mesenchymal transition. Moreover, they observed that LY6E is required for TGF-β signalling and facilitates immune evasion by upregulating PD-L1 on cancer cells, by recruiting T-regulatory cells and dampening NK cell activation (17). In line, our survival analysis of a paediatric ALL-patient cohort confirmed the cancer driving feature of LY6E also in leukaemia. Our data clearly shows that NK cells are less likely to form conjugates with human leukemic cells in the presence of LY6E, which ultimately resulted in a minor increase in NK cell susceptibility. In summary, this substantiates the reported oncogenic and immune suppressive properties of human LY6E and murine Ly6A, but the exact mechanism remains unknown. Although Ly6A has not been directly associated with cell metabolism, our data indicates that *Ly6a* KO B-ALL cells produce more ATP through glycolysis and oxidative phosphorylation. Increased metabolism is generally associated with immune suppression and NK cell inhibition (76), which is contradictory to our observed phenotype. However, it was also shown that increased glycolysis can enhance the expression of NK cell activating NKG2D ligands (51). We only observed a slight and insignificant increase of Rae1 on *Ly6a* KO B-ALL cells, besides differences in CD112 and CD244 expression. It has been suggested that Ly6A regulates the clustering of receptors or ligands within lipid rafts on the cell membrane (77). We thus hypothesise that potential differences in local concentrations of NK cell receptor ligands may contribute to the lower susceptibility of Ly6A-expressing B-ALL cells to NK cell-mediated killing, but this notion needs further evaluation.

Considering the association of *Ly6a* with cancer stemness, our findings align with the concept that immunoedited and thus more aggressive tumour cells may harbour cancer stem cell (CSC) features. CSCs are considered responsible for metastasis and therapy resistance, as they are characterised by deregulated differentiation, self-renewal, drug resistance, and amongst others evasion of immunosurveillance. Unlike other immune cells, NK cells are believed to show direct cytotoxicity against CSCs (78). On the other hand, as CSCs frequently upregulate classical and non-classical MHC-I molecules while losing NK cell activating ligands, it is still a matter of debate if and how NK cells are able to detect and kill CSCs (78,79).

In conclusion, our work sheds light on the intricate interplay between NK and leukaemic cells during initial encounters. This study shows that leukaemic cells are actively shaped by NK cells and unravels novel cancer evasion strategies by identifying potential driver genes in mice (*Ly6a*) and men (*LY6E*). Further research, especially *in vivo* and with single cell resolution, is required to delve deeper into primary and secondary resistance mechanisms and the general role of Ly6A and LY6E in NK cell-mediated tumour surveillance. These findings ultimately deepen our understanding of the initial interaction between tumour and NK cells and of cancer immune evasion in general.

## Acknowledgements

We thank Veronika Sexl (VetMedUni Vienna, Austria) for providing the BCR/ABL^p185^ construct, Thomas Kolbe (VetMedUni Vienna, Austria) for providing the *B2m^−/−^*mice and Sabine Strehl and Klaus Fortschegger (CCRI, Austria) for discussions and sharing information. We are grateful to Marcus Toetzl for assistance with ATAC-seq library preparation and Melanie Hofmann and Gernot Schabbauer for support with the Seahorse experiments. We thank the CCRI FACS and Bioinformatics Core Units for assistance with cell sorting and bioinformatics/statistics, respectively, the Core Facility Laboratory Animal Breeding and Husbandry of the Medical University of Vienna (MUW, Austria) for taking care of our mice and the Biomedical Sequencing Facility and the Proteomics Facility of the CeMM Research Centre for Molecular Medicine of the Austrian Academy of Sciences for their support on next-generation sequencing and mass spectrometry analysis, respectively. This work was supported by the St. Anna Children’s Cancer Research Institute (Vienna, Austria), by a DOC fellowship of the Austrian Academy of Sciences awarded to M.C.B. (#25905), and by grants from the Fellinger Krebsforschung to E.M.P. and from the Austrian Science Fund (10.55776/P32001-B, 10.55776/P34832 and 10.55776/WKP132-B to E.M.P.; 10.55776/TAI 454 and 10.55776/TAI 732 to F.H.; 10.55776/P31563 and 10.55776/I4649 to X.K.). F.H. was supported by funding from the Alex’s Lemonade Stand Foundation for Childhood Cancer (20–17258) and A.R.P. was supported by the Mildred Scheel Early Career Center Dresden P2, funded by the German Cancer Aid.

## Author contributions

Conceptualisatìon: E.M.P.

Methodology: M.C.B., B.K., E.M.P., with input from J.U., J.C., L.P., T.S., D.G.,

Formal analysis: M.C.B., M.R.S, A.B., P.R., D.S., A.R.P., E.M.P.

Investigation: M.C.B., H.B., A.D., F.O.D., B.K., F.H.-G., S.D., L.S., S.S., D.E., D.G.

Writing (original draft): M.C.B., E.M.P.

Visualisation: M.C.B., M.R.S., A.B., X.K., E.M.P.

Resources: J.U., J.C., M.L., L.P., T.S., F.H., E.M.P.

Supervision: F.H., E.M.P.

Funding acquisition: M.C.B., E.M.P.

All authors reviewed and edited the manuscript.

