## Supplementary Table 1 for "Natural killer cell cytotoxicity shapes the clonal evolution of B cell leukaemia"

**Table S1: Primers**

| Name | Sequence | Usage | Annealing Temp. in °C | Polymerase |
|---|---|---|---|---|
| BC1DS_lib_oligo | AAGCTTTTGCTGCCGTCAACTAGAACACTCGAGATCAGNNNNNNNNNNNNNNNNNNNNNATGTGGTATGATGTATCATCTGAGTAGCTAGTTCCGCAACTACCTTCAGGGACCTGGTGCTGTTCATCCTACTCTTGATCTTGAATTCATAGATATCGTCTGTACTCAGTTGTCACTTTCAT | Barcode library generation | 57 | Phusion polymerase (New England Biolabs (NEB)) |
| BC1DS_lib_rev_AscI | ATGAAAGTGACAACTGAGTACAGACGGCGCGCCTATGAATTCAAGATCAAGAG | Barcode library generation | 57 | Phusion polymerase (NEB) |
| topLIB | TGCTGCCGTCAACTAGAACA | Barcode library preparation PCR#1 | 63.7 | NEBNext Ultra II Q5 Master Mix (NEB) |
| botLIB | GATCTCGAATCAGGCGCTTA | Barcode library preparation PCR#1 | 63.7 | NEBNext Ultra II Q5 Master Mix (NEB) |
| BC1v2DS For | ACACTCTTTCCCTACACGACGCTCTTCCGATCTNNNNCTAGAACACTCGAGATCAG | Barcode library preparation PCR#2 | 63.7 | NEBNext Ultra II Q5 Master Mix (NEB) |
| BC1v2DS Rev | GTGACTGGAGTTCAGACGTGTGCTCTTCCGATCGATCTCGAATCAGGCGCTTA | Barcode library preparation PCR#2 | 63.7 | NEBNext Ultra II Q5 Master Mix (NEB) |
| P5 universal Illumina | AATGATACGGCGACCACCGAGATCTACACTCTTTCCCTACACGACGCTCTTCCGATC*T | Barcode library preparation PCR#3 | 69 | NEBNext Ultra II Q5 Master Mix (NEB) |
| Index P1 | CAAGCAGAAGACGGCATACGAGATGTCGGTAAGTGACTGGAGTTCAGACGTGTGCTCTTCCGATCT | Barcode library preparation PCR#3 | 69 | NEBNext Ultra II Q5 Master Mix (NEB) |
| Index P2 | CAAGCAGAAGACGGCATACGAGATAGGTCACTGTGACTGGAGTTCAGACGTGTGCTCTTCCGATCT | Barcode library preparation PCR#3 | 69 | NEBNext Ultra II Q5 Master Mix (NEB) |
| Index P3 | CAAGCAGAAGACGGCATACGAGATGAATCCGAGTGACTGGAGTTCAGACGTGTGCTCTTCCGATCT | Barcode library preparation PCR#3 | 69 | NEBNext Ultra II Q5 Master Mix (NEB) |
| Index P4 | CAAGCAGAAGACGGCATACGAGATGTACCTTGGTGACTGGAGTTCAGACGTGTGCTCTTCCGATCT | Barcode library preparation PCR#3 | 69 | NEBNext Ultra II Q5 Master Mix (NEB) |
| Index P5 | CAAGCAGAAGACGGCATACGAGATCATGAGGAGTGACTGGAGTTCAGACGTGTGCTCTTCCGATCT | Barcode library preparation PCR#3 | 69 | NEBNext Ultra II Q5 Master Mix (NEB) |
| Index P6 | CAAGCAGAAGACGGCATACGAGATTGACTGACGTGACTGGAGTTCAGACGTGTGCTCTTCCGATCT | Barcode library preparation PCR#3 | 69 | NEBNext Ultra II Q5 Master Mix (NEB) |
| Index P7 | CAAGCAGAAGACGGCATACGAGATCGTATTCGGTGACTGGAGTTCAGACGTGTGCTCTTCCGATCT | Barcode library preparation PCR#3 | 69 | NEBNext Ultra II Q5 Master Mix (NEB) |
| Index P8 | CAAGCAGAAGACGGCATACGAGATCTCCTAGAGTGACTGGAGTTCAGACGTGTGCTCTTCCGATCT | Barcode library preparation PCR#3 | 69 | NEBNext Ultra II Q5 Master Mix (NEB) |
| Index P9 | CAAGCAGAAGACGGCATACGAGATTAGTTGCGGTGACTGGAGTTCAGACGTGTGCTCTTCCGATCT | Barcode library preparation PCR#3 | 69 | NEBNext Ultra II Q5 Master Mix (NEB) |
| Index P10 | CAAGCAGAAGACGGCATACGAGATGAGATACGGTGACTGGAGTTCAGACGTGTGCTCTTCCGATCT | Barcode library preparation PCR#3 | 69 | NEBNext Ultra II Q5 Master Mix (NEB) |
| Index P11 | CAAGCAGAAGACGGCATACGAGATAGGTGTACGTGACTGGAGTTCAGACGTGTGCTCTTCCGATCT | Barcode library preparation PCR#3 | 69 | NEBNext Ultra II Q5 Master Mix (NEB) |
| Index P12 | CAAGCAGAAGACGGCATACGAGATTAATGCCGGTGACTGGAGTTCAGACGTGTGCTCTTCCGATCT | Barcode library preparation PCR#3 | 69 | NEBNext Ultra II Q5 Master Mix (NEB) |
| Index P13 | CAAGCAGAAGACGGCATACGAGATTCAGACGAGTGACTGGAGTTCAGACGTGTGCTCTTCCGATCT | Barcode library preparation PCR#3 | 69 | NEBNext Ultra II Q5 Master Mix (NEB) |
| Index P14 | CAAGCAGAAGACGGCATACGAGATGATAGGCTGTGACTGGAGTTCAGACGTGTGCTCTTCCGATCT | Barcode library preparation PCR#3 | 69 | NEBNext Ultra II Q5 Master Mix (NEB) |
| Index P15 | CAAGCAGAAGACGGCATACGAGATTGGTACAGGTGACTGGAGTTCAGACGTGTGCTCTTCCGATCT | Barcode library preparation PCR#3 | 69 | NEBNext Ultra II Q5 Master Mix (NEB) |
| Index P16 | CAAGCAGAAGACGGCATACGAGATCAAGGTCTGTGACTGGAGTTCAGACGTGTGCTCTTCCGATCT | Barcode library preparation PCR#3 | 69 | NEBNext Ultra II Q5 Master Mix (NEB) |
| Index P17 | CAAGCAGAAGACGGCATACGAGATGCTATCCTGTGACTGGAGTTCAGACGTGTGCTCTTCCGATCT | Barcode library preparation PCR#3 | 69 | NEBNext Ultra II Q5 Master Mix (NEB) |
| Index P18 | CAAGCAGAAGACGGCATACGAGATATGGAAGGGTGACTGGAGTTCAGACGTGTGCTCTTCCGATCT | Barcode library preparation PCR#3 | 69 | NEBNext Ultra II Q5 Master Mix (NEB) |
| Index P19 | CAAGCAGAAGACGGCATACGAGATTCAAGGACGTGACTGGAGTTCAGACGTGTGCTCTTCCGATCT | Barcode library preparation PCR#3 | 69 | NEBNext Ultra II Q5 Master Mix (NEB) |
| Index P20 | CAAGCAGAAGACGGCATACGAGATGTTACGCAGTGACTGGAGTTCAGACGTGTGCTCTTCCGATCT | Barcode library preparation PCR#3 | 69 | NEBNext Ultra II Q5 Master Mix (NEB) |
| Index P21 | CAAGCAGAAGACGGCATACGAGATAGTCTGTGGTGACTGGAGTTCAGACGTGTGCTCTTCCGATCT | Barcode library preparation PCR#3 | 69 | NEBNext Ultra II Q5 Master Mix (NEB) |
| Index P22 | CAAGCAGAAGACGGCATACGAGATGCACGTAAGTGACTGGAGTTCAGACGTGTGCTCTTCCGATCT | Barcode library preparation PCR#3 | 69 | NEBNext Ultra II Q5 Master Mix (NEB) |
| Index P23 | CAAGCAGAAGACGGCATACGAGATAACCTTGGGTGACTGGAGTTCAGACGTGTGCTCTTCCGATCT | Barcode library preparation PCR#3 | 69 | NEBNext Ultra II Q5 Master Mix (NEB) |
| Index P24 | CAAGCAGAAGACGGCATACGAGATATTGCGTGGTGACTGGAGTTCAGACGTGTGCTCTTCCGATCT | Barcode library preparation PCR#3 | 69 | NEBNext Ultra II Q5 Master Mix (NEB) |
| Index P25 | CAAGCAGAAGACGGCATACGAGATACCTGGAAGTGACTGGAGTTCAGACGTGTGCTCTTCCGATCT | Barcode library preparation PCR#3 | 69 | NEBNext Ultra II Q5 Master Mix (NEB) |
| Index P26 | CAAGCAGAAGACGGCATACGAGATGGAGATGAGTGACTGGAGTTCAGACGTGTGCTCTTCCGATCT | Barcode library preparation PCR#3 | 69 | NEBNext Ultra II Q5 Master Mix (NEB) |
| Index P27 | CAAGCAGAAGACGGCATACGAGATGTACTCTCGTGACTGGAGTTCAGACGTGTGCTCTTCCGATCT | Barcode library preparation PCR#3 | 69 | NEBNext Ultra II Q5 Master Mix (NEB) |
| Index P28 | CAAGCAGAAGACGGCATACGAGATGTAACGACGTGACTGGAGTTCAGACGTGTGCTCTTCCGATCT | Barcode library preparation PCR#3 | 69 | NEBNext Ultra II Q5 Master Mix (NEB) |
| Index P29 | CAAGCAGAAGACGGCATACGAGATATTCCTCCGTGACTGGAGTTCAGACGTGTGCTCTTCCGATCT | Barcode library preparation PCR#3 | 69 | NEBNext Ultra II Q5 Master Mix (NEB) |
| Index P30 | CAAGCAGAAGACGGCATACGAGATGTGTTCCTGTGACTGGAGTTCAGACGTGTGCTCTTCCGATCT | Barcode library preparation PCR#3 | 69 | NEBNext Ultra II Q5 Master Mix (NEB) |
| Index P31 | CAAGCAGAAGACGGCATACGAGATAAGCACTGGTGACTGGAGTTCAGACGTGTGCTCTTCCGATCT | Barcode library preparation PCR#3 | 69 | NEBNext Ultra II Q5 Master Mix (NEB) |
| Index P32 | CAAGCAGAAGACGGCATACGAGATCTAGCAAGGTGACTGGAGTTCAGACGTGTGCTCTTCCGATCT | Barcode library preparation PCR#3 | 69 | NEBNext Ultra II Q5 Master Mix (NEB) |
| Index P33 | CAAGCAGAAGACGGCATACGAGATTGCTTCCAGTGACTGGAGTTCAGACGTGTGCTCTTCCGATCT | Barcode library preparation PCR#3 | 69 | NEBNext Ultra II Q5 Master Mix (NEB) |
| Index P34 | CAAGCAGAAGACGGCATACGAGATGCTTAGCTGTGACTGGAGTTCAGACGTGTGCTCTTCCGATCT | Barcode library preparation PCR#3 | 69 | NEBNext Ultra II Q5 Master Mix (NEB) |
| Index P35 | CAAGCAGAAGACGGCATACGAGATAACCGTTCGTGACTGGAGTTCAGACGTGTGCTCTTCCGATCT | Barcode library preparation PCR#3 | 69 | NEBNext Ultra II Q5 Master Mix (NEB) |
| Index P36 | CAAGCAGAAGACGGCATACGAGATGACATTCCGTGACTGGAGTTCAGACGTGTGCTCTTCCGATCT | Barcode library preparation PCR#3 | 69 | NEBNext Ultra II Q5 Master Mix (NEB) |
| Index P37 | CAAGCAGAAGACGGCATACGAGATAGACCGTAGTGACTGGAGTTCAGACGTGTGCTCTTCCGATCT | Barcode library preparation PCR#3 | 69 | NEBNext Ultra II Q5 Master Mix (NEB) |
| Index P38 | CAAGCAGAAGACGGCATACGAGATGATACTGGGTGACTGGAGTTCAGACGTGTGCTCTTCCGATCT | Barcode library preparation PCR#3 | 69 | NEBNext Ultra II Q5 Master Mix (NEB) |
| Index P39 | CAAGCAGAAGACGGCATACGAGATTGCGTAGAGTGACTGGAGTTCAGACGTGTGCTCTTCCGATCT | Barcode library preparation PCR#3 | 69 | NEBNext Ultra II Q5 Master Mix (NEB) |
| Index P40 | CAAGCAGAAGACGGCATACGAGATTCGGTTACGTGACTGGAGTTCAGACGTGTGCTCTTCCGATCT | Barcode library preparation PCR#3 | 69 | NEBNext Ultra II Q5 Master Mix (NEB) |
| Index P41 | CAAGCAGAAGACGGCATACGAGATATGACGTCGTGACTGGAGTTCAGACGTGTGCTCTTCCGATCT | Barcode library preparation PCR#3 | 69 | NEBNext Ultra II Q5 Master Mix (NEB) |
| Index P42 | CAAGCAGAAGACGGCATACGAGATGCTGTAAGGTGACTGGAGTTCAGACGTGTGCTCTTCCGATCT | Barcode library preparation PCR#3 | 69 | NEBNext Ultra II Q5 Master Mix (NEB) |
| Index P43 | CAAGCAGAAGACGGCATACGAGATGCAATGGAGTGACTGGAGTTCAGACGTGTGCTCTTCCGATCT | Barcode library preparation PCR#3 | 69 | NEBNext Ultra II Q5 Master Mix (NEB) |
| Index P44 | CAAGCAGAAGACGGCATACGAGATATCTCGCTGTGACTGGAGTTCAGACGTGTGCTCTTCCGATCT | Barcode library preparation PCR#3 | 69 | NEBNext Ultra II Q5 Master Mix (NEB) |
| Index P45 | CAAGCAGAAGACGGCATACGAGATGGCTATTGGTGACTGGAGTTCAGACGTGTGCTCTTCCGATCT | Barcode library preparation PCR#3 | 69 | NEBNext Ultra II Q5 Master Mix (NEB) |
| Index P46 | CAAGCAGAAGACGGCATACGAGATGGTGTCTTGTGACTGGAGTTCAGACGTGTGCTCTTCCGATCT | Barcode library preparation PCR#3 | 69 | NEBNext Ultra II Q5 Master Mix (NEB) |
| Index P47 | CAAGCAGAAGACGGCATACGAGATTCAACTGGGTGACTGGAGTTCAGACGTGTGCTCTTCCGATCT | Barcode library preparation PCR#3 | 69 | NEBNext Ultra II Q5 Master Mix (NEB) |
| Index P48 | CAAGCAGAAGACGGCATACGAGATCTTCACCAGTGACTGGAGTTCAGACGTGTGCTCTTCCGATCT | Barcode library preparation PCR#3 | 69 | NEBNext Ultra II Q5 Master Mix (NEB) |
| Index P49 | CAAGCAGAAGACGGCATACGAGATACGGTCTTGTGACTGGAGTTCAGACGTGTGCTCTTCCGATCT | Barcode library preparation PCR#3 | 69 | NEBNext Ultra II Q5 Master Mix (NEB) |
| Index P50 | CAAGCAGAAGACGGCATACGAGATTCTCGCAAGTGACTGGAGTTCAGACGTGTGCTCTTCCGATCT | Barcode library preparation PCR#3 | 69 | NEBNext Ultra II Q5 Master Mix (NEB) |
| Index P51 | CAAGCAGAAGACGGCATACGAGATGGAATTGCGTGACTGGAGTTCAGACGTGTGCTCTTCCGATCT | Barcode library preparation PCR#3 | 69 | NEBNext Ultra II Q5 Master Mix (NEB) |
| Index P52 | CAAGCAGAAGACGGCATACGAGATACGGATTCGTGACTGGAGTTCAGACGTGTGCTCTTCCGATCT | Barcode library preparation PCR#3 | 69 | NEBNext Ultra II Q5 Master Mix (NEB) |
| Index P53 | CAAGCAGAAGACGGCATACGAGATTTAAGCGGGTGACTGGAGTTCAGACGTGTGCTCTTCCGATCT | Barcode library preparation PCR#3 | 69 | NEBNext Ultra II Q5 Master Mix (NEB) |
| Index P54 | CAAGCAGAAGACGGCATACGAGATTGCAGGTAGTGACTGGAGTTCAGACGTGTGCTCTTCCGATCT | Barcode library preparation PCR#3 | 69 | NEBNext Ultra II Q5 Master Mix (NEB) |
| Index P55 | CAAGCAGAAGACGGCATACGAGATCAATCGACGTGACTGGAGTTCAGACGTGTGCTCTTCCGATCT | Barcode library preparation PCR#3 | 69 | NEBNext Ultra II Q5 Master Mix (NEB) |
| Index P56 | CAAGCAGAAGACGGCATACGAGATGTGCCATAGTGACTGGAGTTCAGACGTGTGCTCTTCCGATCT | Barcode library preparation PCR#3 | 69 | NEBNext Ultra II Q5 Master Mix (NEB) |
| Index P57 | CAAGCAGAAGACGGCATACGAGATTGTTCGAGGTGACTGGAGTTCAGACGTGTGCTCTTCCGATCT | Barcode library preparation PCR#3 | 69 | NEBNext Ultra II Q5 Master Mix (NEB) |
| Index P58 | CAAGCAGAAGACGGCATACGAGATTGGAGTTGGTGACTGGAGTTCAGACGTGTGCTCTTCCGATCT | Barcode library preparation PCR#3 | 69 | NEBNext Ultra II Q5 Master Mix (NEB) |
| Index P59 | CAAGCAGAAGACGGCATACGAGATACGATGACGTGACTGGAGTTCAGACGTGTGCTCTTCCGATCT | Barcode library preparation PCR#3 | 69 | NEBNext Ultra II Q5 Master Mix (NEB) |
| Index P60 | CAAGCAGAAGACGGCATACGAGATTGATGTCCGTGACTGGAGTTCAGACGTGTGCTCTTCCGATCT | Barcode library preparation PCR#3 | 69 | NEBNext Ultra II Q5 Master Mix (NEB) |
| Index P61 | CAAGCAGAAGACGGCATACGAGATTGAACCTGGTGACTGGAGTTCAGACGTGTGCTCTTCCGATCT | Barcode library preparation PCR#3 | 69 | NEBNext Ultra II Q5 Master Mix (NEB) |
| Index P62 | CAAGCAGAAGACGGCATACGAGATCTTCGTTCGTGACTGGAGTTCAGACGTGTGCTCTTCCGATCT | Barcode library preparation PCR#3 | 69 | NEBNext Ultra II Q5 Master Mix (NEB) |
| Index P63 | CAAGCAGAAGACGGCATACGAGATCTTCTGAGGTGACTGGAGTTCAGACGTGTGCTCTTCCGATCT | Barcode library preparation PCR#3 | 69 | NEBNext Ultra II Q5 Master Mix (NEB) |
| Index P64 | CAAGCAGAAGACGGCATACGAGATTGCTCATGGTGACTGGAGTTCAGACGTGTGCTCTTCCGATCT | Barcode library preparation PCR#3 | 69 | NEBNext Ultra II Q5 Master Mix (NEB) |
| Index P65 | CAAGCAGAAGACGGCATACGAGATAGTTCGTCGTGACTGGAGTTCAGACGTGTGCTCTTCCGATCT | Barcode library preparation PCR#3 | 69 | NEBNext Ultra II Q5 Master Mix (NEB) |
| Index P66 | CAAGCAGAAGACGGCATACGAGATTAGCGTCTGTGACTGGAGTTCAGACGTGTGCTCTTCCGATCT | Barcode library preparation PCR#3 | 69 | NEBNext Ultra II Q5 Master Mix (NEB) |
| Index P67 | CAAGCAGAAGACGGCATACGAGATGGCGTTATGTGACTGGAGTTCAGACGTGTGCTCTTCCGATCT | Barcode library preparation PCR#3 | 69 | NEBNext Ultra II Q5 Master Mix (NEB) |
| Index P68 | CAAGCAGAAGACGGCATACGAGATGGTGATTCGTGACTGGAGTTCAGACGTGTGCTCTTCCGATCT | Barcode library preparation PCR#3 | 69 | NEBNext Ultra II Q5 Master Mix (NEB) |
| Index P69 | CAAGCAGAAGACGGCATACGAGATAACTTGCCGTGACTGGAGTTCAGACGTGTGCTCTTCCGATCT | Barcode library preparation PCR#3 | 69 | NEBNext Ultra II Q5 Master Mix (NEB) |
| Index P70 | CAAGCAGAAGACGGCATACGAGATGCAAGATCGTGACTGGAGTTCAGACGTGTGCTCTTCCGATCT | Barcode library preparation PCR#3 | 69 | NEBNext Ultra II Q5 Master Mix (NEB) |
| Index P71 | CAAGCAGAAGACGGCATACGAGATTCGCATTGGTGACTGGAGTTCAGACGTGTGCTCTTCCGATCT | Barcode library preparation PCR#3 | 69 | NEBNext Ultra II Q5 Master Mix (NEB) |
| Index P72 | CAAGCAGAAGACGGCATACGAGATTGTACACCGTGACTGGAGTTCAGACGTGTGCTCTTCCGATCT | Barcode library preparation PCR#3 | 69 | NEBNext Ultra II Q5 Master Mix (NEB) |
| Index P73 | CAAGCAGAAGACGGCATACGAGATAGCTCCTAGTGACTGGAGTTCAGACGTGTGCTCTTCCGATCT | Barcode library preparation PCR#3 | 69 | NEBNext Ultra II Q5 Master Mix (NEB) |
| Index P74 | CAAGCAGAAGACGGCATACGAGATGCAATTCGGTGACTGGAGTTCAGACGTGTGCTCTTCCGATCT | Barcode library preparation PCR#3 | 69 | NEBNext Ultra II Q5 Master Mix (NEB) |
| Index P75 | CAAGCAGAAGACGGCATACGAGATCTTAGGACGTGACTGGAGTTCAGACGTGTGCTCTTCCGATCT | Barcode library preparation PCR#3 | 69 | NEBNext Ultra II Q5 Master Mix (NEB) |
| Index P76 | CAAGCAGAAGACGGCATACGAGATGTCCTAAGGTGACTGGAGTTCAGACGTGTGCTCTTCCGATCT | Barcode library preparation PCR#3 | 69 | NEBNext Ultra II Q5 Master Mix (NEB) |
| Index P77 | CAAGCAGAAGACGGCATACGAGATAACGTGGAGTGACTGGAGTTCAGACGTGTGCTCTTCCGATCT | Barcode library preparation PCR#3 | 69 | NEBNext Ultra II Q5 Master Mix (NEB) |
| Index P78 | CAAGCAGAAGACGGCATACGAGATCTGTTGGTGACTGGAGTTCAGACGTGTGCTCTTCCGATCT | Barcode library preparation PCR#3 | 69 | NEBNext Ultra II Q5 Master Mix (NEB) |
| Index P79 | CAAGCAGAAGACGGCATACGAGATGTTAAGGCGTGACTGGAGTTCAGACGTGTGCTCTTCCGATCT | Barcode library preparation PCR#3 | 69 | NEBNext Ultra II Q5 Master Mix (NEB) |
| Index P80 | CAAGCAGAAGACGGCATACGAGATCACCTTACGTGACTGGAGTTCAGACGTGTGCTCTTCCGATCT | Barcode library preparation PCR#3 | 69 | NEBNext Ultra II Q5 Master Mix (NEB) |
| Index P81 | CAAGCAGAAGACGGCATACGAGATTGGTAGCTGTGACTGGAGTTCAGACGTGTGCTCTTCCGATCT | Barcode library preparation PCR#3 | 69 | NEBNext Ultra II Q5 Master Mix (NEB) |
| Index P82 | CAAGCAGAAGACGGCATACGAGATCAGTGAAGGTGACTGGAGTTCAGACGTGTGCTCTTCCGATCT | Barcode library preparation PCR#3 | 69 | NEBNext Ultra II Q5 Master Mix (NEB) |
| Index P83 | CAAGCAGAAGACGGCATACGAGATGTTCAACCGTGACTGGAGTTCAGACGTGTGCTCTTCCGATCT | Barcode library preparation PCR#3 | 69 | NEBNext Ultra II Q5 Master Mix (NEB) |
| Index P84 | CAAGCAGAAGACGGCATACGAGATTGGCTATCGTGACTGGAGTTCAGACGTGTGCTCTTCCGATCT | Barcode library preparation PCR#3 | 69 | NEBNext Ultra II Q5 Master Mix (NEB) |
| Index P85 | CAAGCAGAAGACGGCATACGAGATCTGGAGTAGTGACTGGAGTTCAGACGTGTGCTCTTCCGATCT | Barcode library preparation PCR#3 | 69 | NEBNext Ultra II Q5 Master Mix (NEB) |
| Index P86 | CAAGCAGAAGACGGCATACGAGATTCTCTTCCGTGACTGGAGTTCAGACGTGTGCTCTTCCGATCT | Barcode library preparation PCR#3 | 69 | NEBNext Ultra II Q5 Master Mix (NEB) |
| Index P87 | CAAGCAGAAGACGGCATACGAGATTCTAACGCGTGACTGGAGTTCAGACGTGTGCTCTTCCGATCT | Barcode library preparation PCR#3 | 69 | NEBNext Ultra II Q5 Master Mix (NEB) |
| Index P88 | CAAGCAGAAGACGGCATACGAGATGGTCAGATGTGACTGGAGTTCAGACGTGTGCTCTTCCGATCT | Barcode library preparation PCR#3 | 69 | NEBNext Ultra II Q5 Master Mix (NEB) |
| Index P89 | CAAGCAGAAGACGGCATACGAGATCTCTGGTTGTGACTGGAGTTCAGACGTGTGCTCTTCCGATCT | Barcode library preparation PCR#3 | 69 | NEBNext Ultra II Q5 Master Mix (NEB) |
| Index P90 | CAAGCAGAAGACGGCATACGAGATTGTGGTACGTGACTGGAGTTCAGACGTGTGCTCTTCCGATCT | Barcode library preparation PCR#3 | 69 | NEBNext Ultra II Q5 Master Mix (NEB) |
| Index P91 | CAAGCAGAAGACGGCATACGAGATCCTATACCGTGACTGGAGTTCAGACGTGTGCTCTTCCGATCT | Barcode library preparation PCR#3 | 69 | NEBNext Ultra II Q5 Master Mix (NEB) |
| Index P92 | CAAGCAGAAGACGGCATACGAGATTTCTCTCGGTGACTGGAGTTCAGACGTGTGCTCTTCCGATCT | Barcode library preparation PCR#3 | 69 | NEBNext Ultra II Q5 Master Mix (NEB) |
| Index P93 | CAAGCAGAAGACGGCATACGAGATGTATGCTGGTGACTGGAGTTCAGACGTGTGCTCTTCCGATCT | Barcode library preparation PCR#3 | 69 | NEBNext Ultra II Q5 Master Mix (NEB) |
| Index P94 | CAAGCAGAAGACGGCATACGAGATAAGTCGAGGTGACTGGAGTTCAGACGTGTGCTCTTCCGATCT | Barcode library preparation PCR#3 | 69 | NEBNext Ultra II Q5 Master Mix (NEB) |
| Index P95 | CAAGCAGAAGACGGCATACGAGATAACCGAAGGTGACTGGAGTTCAGACGTGTGCTCTTCCGATCT | Barcode library preparation PCR#3 | 69 | NEBNext Ultra II Q5 Master Mix (NEB) |
| Index P96 | CAAGCAGAAGACGGCATACGAGATTGTTGTGGGTGACTGGAGTTCAGACGTGTGCTCTTCCGATCT | Barcode library preparation PCR#3 | 69 | NEBNext Ultra II Q5 Master Mix (NEB) |
| Ad1_noMX | AATGATACGGCGACCACCGAGATCTACACTCGTCGGCAGCGTCAGATGTG | ATAC sample library preparation P5 adapter | 63 | NEBNext Ultra II Q5 Master Mix (NEB) |
| Ad2.1 | CAAGCAGAAGACGGCATACGAGATTCGCCTTAGTCTCGTGGGCTCGGAGATGT | ATAC sample library preparation P7 adapter and index | 63 | NEBNext Ultra II Q5 Master Mix (NEB) |
| Ad2.2 | CAAGCAGAAGACGGCATACGAGATCTAGTACGGTCTCGTGGGCTCGGAGATGT | ATAC sample library preparation P7 adapter and index | 63 | NEBNext Ultra II Q5 Master Mix (NEB) |
| Ad2.3 | CAAGCAGAAGACGGCATACGAGATTTCTGCCTGTCTCGTGGGCTCGGAGATGT | ATAC sample library preparation P7 adapter and index | 63 | NEBNext Ultra II Q5 Master Mix (NEB) |
| Ad2.4 | CAAGCAGAAGACGGCATACGAGATGCTCAGGAGTCTCGTGGGCTCGGAGATGT | ATAC sample library preparation P7 adapter and index | 63 | NEBNext Ultra II Q5 Master Mix (NEB) |
| Ad2.5 | CAAGCAGAAGACGGCATACGAGATAGGAGTCCGTCTCGTGGGCTCGGAGATGT | ATAC sample library preparation P7 adapter and index | 63 | NEBNext Ultra II Q5 Master Mix (NEB) |
| Ad2.6 | CAAGCAGAAGACGGCATACGAGATCATGCCTAGTCTCGTGGGCTCGGAGATGT | ATAC sample library preparation P7 adapter and index | 63 | NEBNext Ultra II Q5 Master Mix (NEB) |
| Ad2.7 | CAAGCAGAAGACGGCATACGAGATGTAGAGAGGTCTCGTGGGCTCGGAGATGT | ATAC sample library preparation P7 adapter and index | 63 | NEBNext Ultra II Q5 Master Mix (NEB) |
| Ad2.8 | CAAGCAGAAGACGGCATACGAGATCCTCTCTGGTCTCGTGGGCTCGGAGATGT | ATAC sample library preparation P7 adapter and index | 63 | NEBNext Ultra II Q5 Master Mix (NEB) |
| Ad2.9 | CAAGCAGAAGACGGCATACGAGATAGCGTAGCGTCTCGTGGGCTCGGAGATGT | ATAC sample library preparation P7 adapter and index | 63 | NEBNext Ultra II Q5 Master Mix (NEB) |
| Ad2.10 | CAAGCAGAAGACGGCATACGAGATCAGCCTCGGTCTCGTGGGCTCGGAGATGT | ATAC sample library preparation P7 adapter and index | 63 | NEBNext Ultra II Q5 Master Mix (NEB) |
| Ad2.11 | CAAGCAGAAGACGGCATACGAGATTGCCTCTTGTCTCGTGGGCTCGGAGATGT | ATAC sample library preparation P7 adapter and index | 63 | NEBNext Ultra II Q5 Master Mix (NEB) |
| Ad2.12 | CAAGCAGAAGACGGCATACGAGATTCCTCTACGTCTCGTGGGCTCGGAGATGT | ATAC sample library preparation P7 adapter and index | 63 | NEBNext Ultra II Q5 Master Mix (NEB) |
| Ad2.13 | CAAGCAGAAGACGGCATACGAGATATCACGACGTCTCGTGGGCTCGGAGATGT | ATAC sample library preparation P7 adapter and index | 63 | NEBNext Ultra II Q5 Master Mix (NEB) |
| Ad2.14 | CAAGCAGAAGACGGCATACGAGATACAGTGGTGTCTCGTGGGCTCGGAGATGT | ATAC sample library preparation P7 adapter and index | 63 | NEBNext Ultra II Q5 Master Mix (NEB) |
| Ad2.15 | CAAGCAGAAGACGGCATACGAGATCAGATCCAGTCTCGTGGGCTCGGAGATGT | ATAC sample library preparation P7 adapter and index | 63 | NEBNext Ultra II Q5 Master Mix (NEB) |
| Ad2.16 | CAAGCAGAAGACGGCATACGAGATACAAACGGGTCTCGTGGGCTCGGAGATGT | ATAC sample library preparation P7 adapter and index | 63 | NEBNext Ultra II Q5 Master Mix (NEB) |
| Ad2.17 | CAAGCAGAAGACGGCATACGAGATACCCAGCAGTCTCGTGGGCTCGGAGATGT | ATAC sample library preparation P7 adapter and index | 63 | NEBNext Ultra II Q5 Master Mix (NEB) |
| Ad2.18 | CAAGCAGAAGACGGCATACGAGATAACCCCTCGTCTCGTGGGCTCGGAGATGT | ATAC sample library preparation P7 adapter and index | 63 | NEBNext Ultra II Q5 Master Mix (NEB) |
| Ad2.19 | CAAGCAGAAGACGGCATACGAGATCCCAACCTGTCTCGTGGGCTCGGAGATGT | ATAC sample library preparation P7 adapter and index | 63 | NEBNext Ultra II Q5 Master Mix (NEB) |
| Ad2.20 | CAAGCAGAAGACGGCATACGAGATCACCACACGTCTCGTGGGCTCGGAGATGT | ATAC sample library preparation P7 adapter and index | 63 | NEBNext Ultra II Q5 Master Mix (NEB) |
| Ad2.21 | CAAGCAGAAGACGGCATACGAGATGAAACCCAGTCTCGTGGGCTCGGAGATGT | ATAC sample library preparation P7 adapter and index | 63 | NEBNext Ultra II Q5 Master Mix (NEB) |
| Ad2.22 | CAAGCAGAAGACGGCATACGAGATTGTGACCAGTCTCGTGGGCTCGGAGATGT | ATAC sample library preparation P7 adapter and index | 63 | NEBNext Ultra II Q5 Master Mix (NEB) |
| Ad2.23 | CAAGCAGAAGACGGCATACGAGATAGGGTCAAGTCTCGTGGGCTCGGAGATGT | ATAC sample library preparation P7 adapter and index | 63 | NEBNext Ultra II Q5 Master Mix (NEB) |
| Ad2.24 | CAAGCAGAAGACGGCATACGAGATAGGAGTGGGTCTCGTGGGCTCGGAGATGT | ATAC sample library preparation P7 adapter and index | 63 | NEBNext Ultra II Q5 Master Mix (NEB) |
| Ad2.25 | CAAGCAGAAGACGGCATACGAGATCACTTTGTGTCTCGTGGGCTCGGAGATGT | ATAC sample library preparation P7 adapter and index | 63 | NEBNext Ultra II Q5 Master Mix (NEB) |
| Ad2.26 | CAAGCAGAAGACGGCATACGAGATTTCAAGTAGTCTCGTGGGCTCGGAGATGT | ATAC sample library preparation P7 adapter and index | 63 | NEBNext Ultra II Q5 Master Mix (NEB) |
| Ad2.27 | CAAGCAGAAGACGGCATACGAGATGCTATCACGTCTCGTGGGCTCGGAGATGT | ATAC sample library preparation P7 adapter and index | 63 | NEBNext Ultra II Q5 Master Mix (NEB) |
| Ad2.28 | CAAGCAGAAGACGGCATACGAGATAATCTACTGTCTCGTGGGCTCGGAGATGT | ATAC sample library preparation P7 adapter and index | 63 | NEBNext Ultra II Q5 Master Mix (NEB) |
| Ad2.29 | CAAGCAGAAGACGGCATACGAGATCCGGCAATGTCTCGTGGGCTCGGAGATGT | ATAC sample library preparation P7 adapter and index | 63 | NEBNext Ultra II Q5 Master Mix (NEB) |
| Ad2.30 | CAAGCAGAAGACGGCATACGAGATCTTAGCAAGTCTCGTGGGCTCGGAGATGT | ATAC sample library preparation P7 adapter and index | 63 | NEBNext Ultra II Q5 Master Mix (NEB) |
| Ad2.31 | CAAGCAGAAGACGGCATACGAGATTAACTTATGTCTCGTGGGCTCGGAGATGT | ATAC sample library preparation P7 adapter and index | 63 | NEBNext Ultra II Q5 Master Mix (NEB) |
| Ad2.32 | CAAGCAGAAGACGGCATACGAGATCGAGTGATGTCTCGTGGGCTCGGAGATGT | ATAC sample library preparation P7 adapter and index | 63 | NEBNext Ultra II Q5 Master Mix (NEB) |
| Ad2.33 | CAAGCAGAAGACGGCATACGAGATCTGTTAACGTCTCGTGGGCTCGGAGATGT | ATAC sample library preparation P7 adapter and index | 63 | NEBNext Ultra II Q5 Master Mix (NEB) |
| Ad2.34 | CAAGCAGAAGACGGCATACGAGATCTACCATTGTCTCGTGGGCTCGGAGATGT | ATAC sample library preparation P7 adapter and index | 63 | NEBNext Ultra II Q5 Master Mix (NEB) |
| Ad2.35 | CAAGCAGAAGACGGCATACGAGATACGTGCTCGTCTCGTGGGCTCGGAGATGT | ATAC sample library preparation P7 adapter and index | 63 | NEBNext Ultra II Q5 Master Mix (NEB) |
| Ad2.36 | CAAGCAGAAGACGGCATACGAGATTGACGAAAGTCTCGTGGGCTCGGAGATGT | ATAC sample library preparation P7 adapter and index | 63 | NEBNext Ultra II Q5 Master Mix (NEB) |
| Ad2.37 | CAAGCAGAAGACGGCATACGAGATAATTCTTGGTCTCGTGGGCTCGGAGATGT | ATAC sample library preparation P7 adapter and index | 63 | NEBNext Ultra II Q5 Master Mix (NEB) |
| Ad2.38 | CAAGCAGAAGACGGCATACGAGATGGCATTTCGTCTCGTGGGCTCGGAGATGT | ATAC sample library preparation P7 adapter and index | 63 | NEBNext Ultra II Q5 Master Mix (NEB) |
| Ad2.39 | CAAGCAGAAGACGGCATACGAGATATGGCGTTGTCTCGTGGGCTCGGAGATGT | ATAC sample library preparation P7 adapter and index | 63 | NEBNext Ultra II Q5 Master Mix (NEB) |
| Ad2.40 | CAAGCAGAAGACGGCATACGAGATCTGCGAGGGTCTCGTGGGCTCGGAGATGT | ATAC sample library preparation P7 adapter and index | 63 | NEBNext Ultra II Q5 Master Mix (NEB) |
| Ad2.41 | CAAGCAGAAGACGGCATACGAGATGAGGTGTAGTCTCGTGGGCTCGGAGATGT | ATAC sample library preparation P7 adapter and index | 63 | NEBNext Ultra II Q5 Master Mix (NEB) |
| Ad2.42 | CAAGCAGAAGACGGCATACGAGATAAATGACCGTCTCGTGGGCTCGGAGATGT | ATAC sample library preparation P7 adapter and index | 63 | NEBNext Ultra II Q5 Master Mix (NEB) |
| Ad2.43 | CAAGCAGAAGACGGCATACGAGATTAAGATTGGTCTCGTGGGCTCGGAGATGT | ATAC sample library preparation P7 adapter and index | 63 | NEBNext Ultra II Q5 Master Mix (NEB) |
| Ad2.44 | CAAGCAGAAGACGGCATACGAGATAAGGCACAGTCTCGTGGGCTCGGAGATGT | ATAC sample library preparation P7 adapter and index | 63 | NEBNext Ultra II Q5 Master Mix (NEB) |
| Ad2.45 | CAAGCAGAAGACGGCATACGAGATTAATAAGAGTCTCGTGGGCTCGGAGATGT | ATAC sample library preparation P7 adapter and index | 63 | NEBNext Ultra II Q5 Master Mix (NEB) |
| Ad2.46 | CAAGCAGAAGACGGCATACGAGATACTAAGTCGTCTCGTGGGCTCGGAGATGT | ATAC sample library preparation P7 adapter and index | 63 | NEBNext Ultra II Q5 Master Mix (NEB) |
| Ad2.47 | CAAGCAGAAGACGGCATACGAGATGCTGGTCTGTCTCGTGGGCTCGGAGATGT | ATAC sample library preparation P7 adapter and index | 63 | NEBNext Ultra II Q5 Master Mix (NEB) |
| Ad2.48 | CAAGCAGAAGACGGCATACGAGATCTGTATTTGTCTCGTGGGCTCGGAGATGT | ATAC sample library preparation P7 adapter and index | 63 | NEBNext Ultra II Q5 Master Mix (NEB) |
| Ly6a_TIDE_FW | CCTGGGCTTCTGTAAGAGCA | TIDE PCR for Ly6a knockout verification | 68 | NEBNext Ultra II Q5 Master Mix (NEB) |
| Ly6a_TIDE_RV | CAGCAGGGGCTATAAAGGCA | TIDE PCR for Ly6a knockout verification | 68 | NEBNext Ultra II Q5 Master Mix (NEB) |
| Plaat3_TIDE1_FW | TTTGTGAGGTGAGCCAGCAT | TIDE PCR for Plaat3 knockout verification | 68 | NEBNext Ultra II Q5 Master Mix (NEB) |
| Plaat3_TIDE1_RV | CTCTGTGTGGAGCCTGGAAG | TIDE PCR for Plaat3 knockout verification | 68 | NEBNext Ultra II Q5 Master Mix (NEB) |
| Plaat3_TIDE2_FW | ACTGTGTGCATCTAACCCGT | TIDE PCR for Plaat3 knockout verification | 68 | NEBNext Ultra II Q5 Master Mix (NEB) |
| Plaat3_TIDE2_RV | AATGATCGTGACCTGTCCTGG | TIDE PCR for Plaat3 knockout verification | 68 | NEBNext Ultra II Q5 Master Mix (NEB) |
| LY6E_TIDE_FW | GGGGTGTCACTGTCTTTGCT | TIDE PCR for LY6E knockout verification | 64.1 | Q5 Hot Start High Fidelity 2X Master Mix (NEB) |
| LY6E_TIDE_RV | CAGATCAGGCACGTGAGGAG | TIDE PCR for LY6E knockout verification | 64.1 | Q5 Hot Start High Fidelity 2X Master Mix (NEB) |
| qPCR mGapdh_FW | TGTGTCCGTCGTGGATCTGA | qPCR primer mouse housekeeping gene | 60 | Luna universal qPCR Master Mix (NEB) |
| qPCR mGapdh_RV | CCTGCTTCACCACCTTCTTGA | qPCR primer mouse housekeeping gene | 60 | Luna universal qPCR Master Mix (NEB) |
| qPCR mPlaat3_FW | TGCCTACGCGAGAACTGTGA | qPCR primer mPlaat3 | 60 | Luna universal qPCR Master Mix (NEB) |
| qPCR mPlaat3_RV | TACCGCCTTGACCGCATCTC | qPCR primer mPlaat3 | 60 | Luna universal qPCR Master Mix (NEB) |
