## Supplementary Table 2 for "Natural killer cell cytotoxicity shapes the clonal evolution of B cell leukaemia"

| Table S2: Antibodies and viability dyes used for flow cytometry and immunoblotting | | | | | | | |
|---|---|---|---|---|---|---|---|
| **Method** | **Species reactivity** | **Antigen** | **Conjugate** | **Clone** | **Supplier** | **Cat#** | **working dilution** |
| FACS | mouse | CD112 (Nectin-2) | BV786 | 829038 | Becton Dickinson | 748050 | 1/100 |
| FACS | mouse | CD137L (4-1BBL) | PE-Vio615 | REA962 | Miltenyi Biotec | 130-116-091 | 1/100 |
| FACS | mouse | CD154 (CD40L) | APC | REA785 | Miltenyi Biotec | 130-111-361 | 1/100 |
| FACS | mouse | CD155 (PVR) | VioBright FITC | REA519 | Miltenyi Biotec | 130-108-082 | 1/20 |
| FACS | human | CD16 (FcgRIII) | VioGreen | REA423 | Miltenyi Biotec | 130-113-959 | 1/50 |
| FACS | mouse | CD16.2 | purified | 3E9 | Biolegend | 149502 | 1/200 |
| FACS | mouse | CD16/32 | purified | 93 | Biolegend | 101302 | 1/200 |
| FACS | mouse | CD161 (NK1.1) | PE-Vio 770 | REA1162 | Miltenyi Biotec | 130-120-509 | 1/400 |
| FACS | mouse | CD19 | APC-Vio770 | REA749 | Miltenyi Biotec | 130-112-038 | 1/800 |
| FACS | mouse | CD19 | PE | 1D3 | BD Biosciences | 09655B | 1/100 |
| FACS | mouse | CD226 (DNAM-1) | BV421 | TX42.1 | Biolegend | 133615 | 1/100 |
| FACS | mouse | CD262 (TRAIL-R2) | PE | REA1233 | Miltenyi Biotec | 130-124-480 | 1/100 |
| FACS | mouse | CD244.2 (2B4) | PE | 2B4 | BD Pharmingen | 553306 | 1/50 |
| FACS | human | CD3 | PerCP-Vio700 | REA613 | Miltenyi Biotec | 130-113-703 | 1/100 |
| FACS | mouse | CD3 | APC | REA641 | Miltenyi Biotec | 130-122-943 | 1/200 |
| FACS | mouse | CD335 (NKp46) | FITC | REA815 | Miltenyi Biotec | 130-112-200 | 1/200 |
| FACS | mouse | CD335 (NKp46) | VioBlue | REA815 | Miltenyi Biotec | 130112365 | 1/100 |
| FACS | human | CD45 | VioBlue | REA747 | Miltenyi Biotec | 130-110-637 | 1/100 |
| FACS | mouse | CD45.2 | PerCP-Vio700 | REA1223 | Miltenyi Biotec | 130-124-087 | 1/400 |
| FACS | mouse | CD45.2 | VioGreen | REA1223 | Miltenyi Biotec | 130-124-083 | 1/200 |
| FACS | mouse | CD48 | VioBlue | REA1238 | Miltenyi Biotec | 130-124-728 | 1/100 |
| FACS | human | CD56 | PE | REA196 | Miltenyi Biotec | 130-113-312 | 1/400 |
| FACS | mouse | CD69 | APC-Vio770 | REA937 | Miltenyi Biotec | 130-115-578 | 1/100 |
| FACS | human/mouse/rat | Granzyme B Antibody, anti-human/mouse/rat, PE, REAfinity™ | PE | REA226 | Miltenyi Biotec | 130-116-486 | 1/100 |
| FACS | mouse | H-2Db | APC-Vio770 | REA619 | Miltenyi Biotec | 130-120-294 | 1/100 |
| FACS | mouse | H-2Kb | PerCP-Vio700 | REA1198 | Miltenyi Biotec | 130-122-848 | 1/100 |
| FACS | human | IFN-g | BV711 | B27 | Becton Dickinson | 564039 | 1/100 |
| FACS | mouse | IFN-g | BV711 | XMG1.2 | Biolegend | 505836 | 1/100 |
| FACS | mouse | Ly6a (Sca-1) | APC | REA422 | Miltenyi Biotec | 130-123-848 | 1/100 |
| FACS | mouse | MULT-1 | PE | 237104 | R&D systems | FAB2588P | 1/100 |
| FACS | mouse | Qa-1b | Biotin | 6A8.6F10.1A6 | Miltenyi Biotec | 130-105-048 | 1/20 |
| FACS | mouse | QA-2 | VioBlue | REA523 | Miltenyi Biotec | 130-107-905 | 1/20 |
| FACS | mouse | Rae-1 | PE-Vio770 | REA723 | Miltenyi Biotec | 130-111-471 | 1/100 |
| FACS | human | TNF-alpha | PE | cA2 | Miltenyi Biotec | 130-120-489 | 1/100 |
| FACS | all | Fixable Viability Dye 440 UV | 440 UV | | Becton Dickinson | 566332 | 1/500 |
| FACS | all | Fixable Viability Dye eF780 | eFluor780 | | Thermo Scientific | 65086518 | 1/1000 |
| FACS | all | Fixable Viability Dye Zombie Red | PE-Texas Red | | Biolegend | 423109 | 1/2000 |
| FACS | all | Streptavidin | BV711 | | Becton Dickinson | 563262 | 1/100 |
| WB | human | LY6E | | Polyclonal | Thermo Scientific | PA5143999 | 1/1000 |
| WB | human/mous | β-Actin | | 13E5 | Cell Signaling | 4970 | 1/1000 |
| WB | rabbit | Goat anti-Rabbit IgG (H+L) Secondary Antibody | DyLight 680 | Polyclonal | Thermo Scientific | 35568 | 1/10000 |
